# Prediction of m6A and m5C at single-molecule resolution reveals a cooccurrence of RNA modifications across the transcriptome

**DOI:** 10.1101/2022.03.14.484124

**Authors:** P. Acera Mateos, A.J. Sethi, A. Ravindran, A. Srivastava, K. Woodward, S. Mahmud, M. Kanchi, M. Guarnacci, J. Xu, Z.W.S. Yuen, Y. Zhou, A. Sneddon, W. Hamilton, J. Gao, L.M. Starrs, R. Hayashi, V. Wickramasinghe, K. Zarnack, T. Preiss, G. Burgio, N. Dehorter, N. Shirokikh, E. Eyras

## Abstract

The epitranscriptome embodies many new and largely unexplored functions of RNA. A significant roadblock hindering progress in epitranscriptomics is the identification of more than one modification in individual transcript molecules. We address this with CHEUI (CH3 (methylation) Estimation Using Ionic current). CHEUI predicts N6-methyladenosine (m6A) and 5-methylcytidine (m5C) in individual molecules from the same sample, the stoichiometry at transcript reference sites, and differential methylation between any two conditions. CHEUI processes observed and expected nanopore direct RNA sequencing signals to achieve high single-molecule, transcript-site, and stoichiometry accuracies in multiple tests using synthetic RNA standards and cell line data. CHEUI’s capability to identify two modification types in the same sample reveals a co-occurrence of m6A and m5C in individual mRNAs in cell line and tissue transcriptomes. CHEUI provides new avenues to discover and study the function of the epitranscriptome.

## INTRODUCTION

The identification of transcriptome-wide maps of two modified ribonucleotides in messenger RNAs (mRNA), 5-methylcytidine (m5C) and N6-methyladenine (m6A)^1–3^, has sparked over the last decade a new and expanding area of epitranscriptomics. Techniques based on immunoprecipitation, enzymatic, or chemical reactivity enrichment methods, coupled with high-throughput sequencing, have uncovered the role of these and other modifications in multiple steps of mRNA metabolism, including translation of mRNA into protein^4,5^, mRNA stability^6^, and mRNA processing such as pre-RNA alternative splicing^7^ and RNA export from the nucleus^8^. Several physiological processes have also been functionally linked with RNA modifications, such as sex determination^9^, early embryonic development^10^, neurogenesis^11^ and learning^12^. Moreover, growing evidence indicates that RNA modification pathways are dysregulated in diseases such as cancer^13^ and neurological disorders^11^. Most of these studies have focused on changes at global or gene levels or on the dysregulation of the RNA modification machinery, whereas little is known about how multiple modifications occur in individual mRNA molecules.

A major roadblock preventing rapid progress in research on RNA modifications is the general lack of universal modification detection methods. Although over 300 naturally occurring RNA modifications have been described^14^, only a handful of them can be mapped and quantified across the transcriptome^15,16^. Nanopore direct RNA sequencing (DRS) is the only currently available technology that can determine the sequence of individual RNA molecules in their native form at a transcriptome-wide level. DRS can capture information about the chemical structure, including naturally occurring covalent modifications in nucleotide residues (nucleotides)^17,18^. Nonetheless, RNA modification detection from DRS signals presents various challenges. The differences between modified and unmodified signals are subtle at single molecule level and depend on the sequence context. Additionally, due to the variable translocation rate of the molecules through the pores and the potential pore-to-pore variability, different copies of the same molecule present considerable signal variations^19^. These challenges necessitate the application of advanced computational models to interpret the signals and identify modification state.

Several computational methods have been developed in the past few years to detect RNA modifications in DRS data. These methods can be broadly grouped into two categories. The first one includes methods that rely on comparing DRS signals between two conditions, one corresponding to a sample of interest, often the wild type (WT) sample, and the other with a reduced presence of a specific modification, usually obtained through a knock-out (KO) or knock-down (KD) of a modification ‘writer’ enzyme or through *in-vitro* transcription. This category includes Nanocompore^20^, Xpore^21^, DRUMMER^22^, nanoDOC^23^, Yanocomp^24^ and Tombo^25^ in *sample comparison* mode, among others, all utilizing the collective properties of DRS signals in the two conditions. This category also includes ELIGOS^26^ and Epinano^27^, which compare base-calling errors between two experiments; and NanoRMS^28^, which compares signal features between two samples. The second category of tools can operate in a single condition, i.e., without using a KO/KD or an otherwise control. This category includes MINES^29^, Nanom6A^30^, and m6Anet^31^, all predicting m6A on specific sequence contexts, Tombo^25^ in *alternate* mode, which identifies transcriptomic sites with potential m5C modification, and Epinano-RMS, which predicts pseudouridine on high stoichiometry sites^28^. Other methods have been recently developed that use one or more of these strategies to predict RNA modifications^32–35^.

Despite the numerous advances in direct RNA modification detection, some major limitations remain. Approaches comparing two conditions generally require a control sample, which can be difficult or impossible to generate. Their modification calling is also indirect, as it relies on changes in the control sample relative to wild type (WT) and these changes may not be directly related to the modification of interest. For instance, depletion of m5C leads to a reduction of hm5C^36^, hence potentially confounding the results. Regarding the methods that use error patterns, they depend on the specific accuracy of the base caller method, which will vary over time with the base caller version and architecture. Moreover, this may not be applicable to all modifications. For instance, it was described that error patterns were not consistent enough to confidently identify m5C methylation^26^. Limitations also exist in methods that work with individual samples. MINES, Nanom6A, and m6Anet only predict m6A modifications in 5′-DR<u>A</u>CH/RR<u>A</u>CH-3′ motifs, and Epinano-RMS only detects pseudouridine in transcriptome sites of high stoichiometry. Additionally, the ability of current methods from both categories to predict stoichiometry is limited. Some of them cannot predict it, whereas others only estimate the stoichiometry at 5′-DR<u>A</u>CH-3′ sites or rely on a control sample devoid of modifications. Transcriptome-wide methods that can predict multiple modifications in individual RNA molecules could enable more precise study of their function.

To advance the field in this direction, we have developed CHEUI (CH3 (methylation) Estimation Using Ionic current) for the prediction of m6A and m5C from the same sample at a transcriptome-wide level in individual molecules, at transcript reference sites, and between conditions. CHEUI is based on a two-stage neural network and was trained using read signals generated from *in-vitro* transcripts (IVTs). We validated CHEUI’s accuracy through a comprehensive set of benchmarking analyses using synthetic RNA standards, orthogonal experiments, and cell line data. Using CHEUI in cell line transcriptomes, we further identified a co-occurrence of m6A and m5C in individual mRNA molecules. CHEUI addresses some of the current limitations in the transcriptome-wide identification of RNA modifications and provides new opportunities for the study of the epitranscriptome.

## RESULTS

### CHEUI detects m6A and m5C in individual reads, transcriptomic sites, and across conditions

For signal preprocessing, CHEUI uses the nanopore read signals corresponding to the 9-mer, composed of five overlapping 5-mers, centered at every single adenosine (A) for m6A or cytosine (C) for m5C (**Fig. 1a**) (**Supp. Fig. 1**). Signal preprocessing further includes derivation of distances between the observed signals and expected unmodified signal values for each 9-mer (**Fig. 1a**) (**Supp. Figs. 2a-2c**). The inclusion of the distance increased accuracy by ∼10% in a test using independent data (**Supp. Fig. 2d**). After preprocessing the signals, CHEUI employs two different modules: CHEUI-solo **(Fig. 1b),** which makes predictions in individual reads and transcript reference sites in given input sample; and CHEUI-diff **(Fig. 1c),** which tests differential methylation between any two samples. CHEUI-solo predicts RNA methylation at two different levels. Model 1 predicts m6A or m5C at nucleotide resolution on individual read signals. Model 2 predicts m6A or m5C at the transcript site level, i.e., relative to a position in the reference transcript, based on the per-read predictions from Model 1 (**Fig. 1b**). Both CHEUI-solo Models 1 and 2 are Convolutional Neural Networks (CNNs) (**Supp. Fig. 3**). CHEUI-diff uses a statistical test to compare the individual read probabilities from CHEUI-solo Model 1 across two conditions, to predict differential stoichiometry of m6A or m5C at each transcriptomic site (**Fig. 1c**). More details about the models are provided in the Methods section.

**Figure 1.**
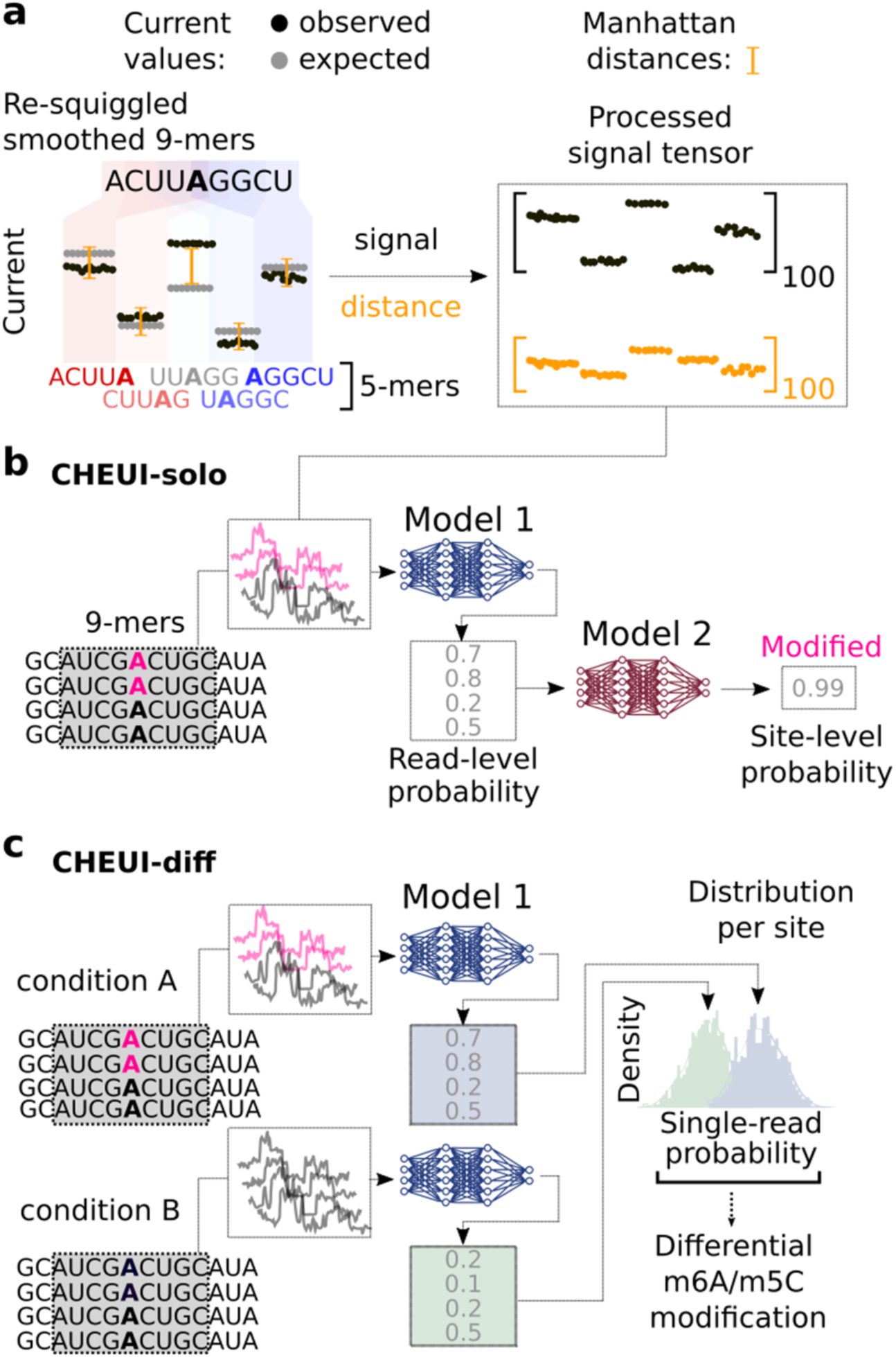
CHEUI architecture, modules, and signal processing approach. **(a)** CHEUI first processes signals for each 9-mer consisting of five consecutive overlapping 5-mers. The signals for each 5-mer are converted into 20 median values, yielding a vector of length 100. A vector of length 100 is obtained for the expected (unmodified) signals for the same five 5-mers and a vector of distances between the expected and observed signal values is calculated. These signal and distance vectors are used as inputs for Model 1. **(b)** CHEUI-solo Model 1 takes the signal and distance vectors corresponding to individual read signals associated with a 9-mer centered at every A (indicated as modified in pink, or unmodified in black) or C, and predicts the probability for each read of being modified A (m6A model) or modified C (m5C model). Model 2 uses the distribution of Model 1 probabilities for all the read signals at each reference transcript site and predicts the probability of the site being methylated and its stoichiometry, which estimated as the proportion of modified reads from Model 1 at that site. **(c)** CHEUI-diff uses the individual read probabilities from Model 1 in any two conditions to test for differential m6A or m5C at reference transcript sites using a Wilcoxon rank-sum test.

### CHEUI accurately detects m5C and m6A in reads and sequence contexts not seen during training

To evaluate CHEUI’s accuracy, we first tested CHEUI-solo’s ability to correctly classify individual read signals not previously used but from 9-mer contexts seen during training, also known as sensor generalization^37^. For this test, only read signals from 9-mers with a single modified nucleotide were considered, i.e., 9-mers where only one A or one C was present, which were collectively called IVT set 1^27^. CHEUI achieved accuracy, precision, and recall values of ∼0.8 for m6A and m5C predictions in individual reads (**Fig. 2a**, IVT set 1). Then, to determine CHEUI’s ability to classify signals from 9-mer contexts not seen during training, also known as *k*-mer generalization^37^, we used signals from a different set of IVTs from a different sequencing experiment^26^, which we called IVT set 2. As before, this test only included signals from 9-mer sites with a single middle A or C. CHEUI achieved accuracy, precision, and recall of ∼0.8 for m6A and ∼0.75 for m5C (**Fig. 2a**, IVT set 2).

**Figure 2.**
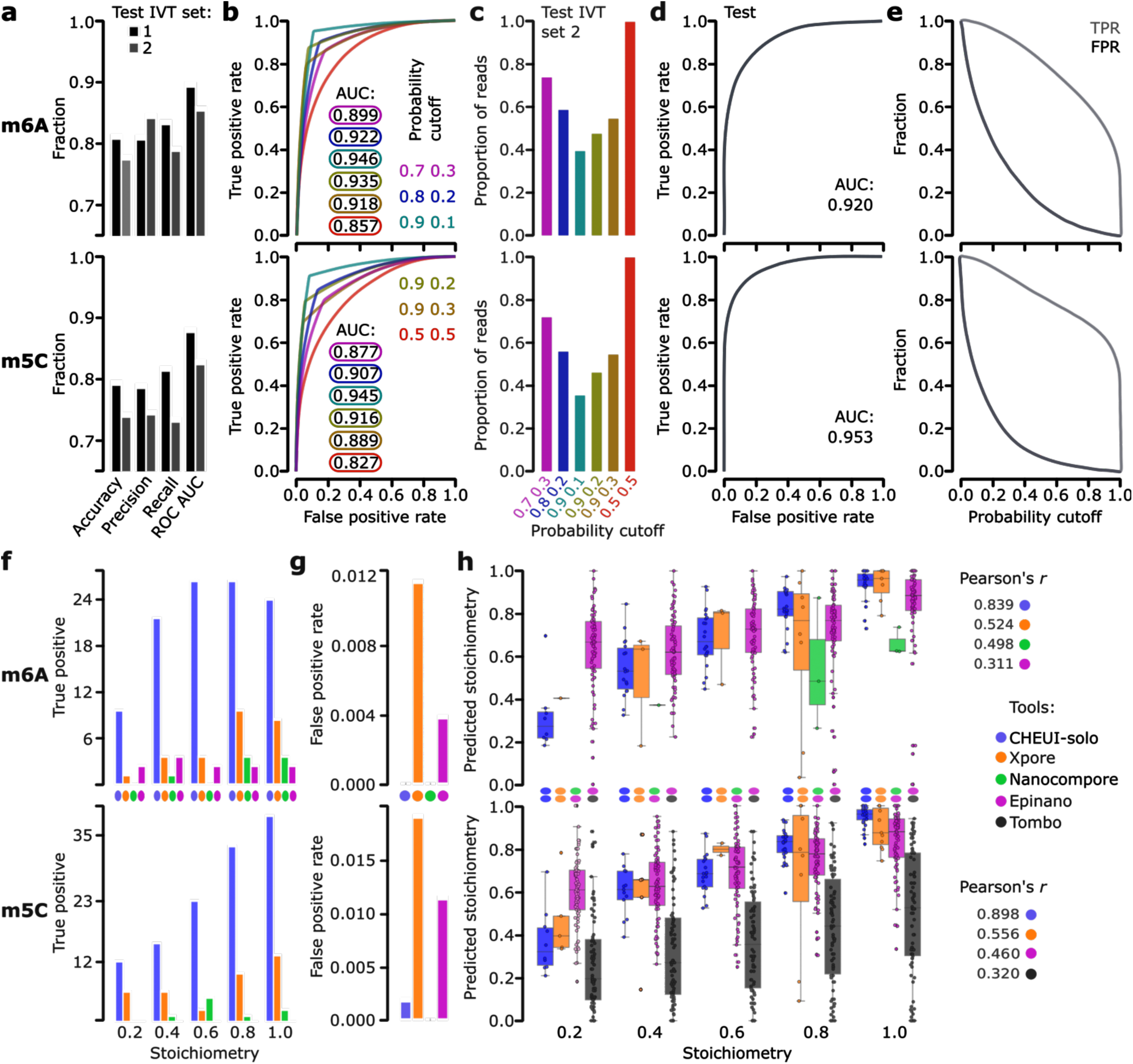
CHEUI’s accuracy metrics at the individual read level and comparison with other RNA modification detection tools. **(a)** Accuracy, precision, recall, and area (AUC) under the receiver operating curve (ROC) for CHEUI-solo Model 1 in m6A (upper panel) and m5C (lower panel) detection. Values are shown for reads containing sequences seen during training (IVT set 1) and for reads with sequences not seen during training (IVT set 2). The metrics (accuracy, precision, recall, AUC) for m6A were (0.835, 0.820, 0.853, 0.910) for IVT set 1 and (0.777, 0.844, 0.791, 0.856) for IVT set 2. For m5C, these metrics were (0.793, 0.788, 0.816, 0.879) for IVT set 1 and (0.741, 0.745, 0.733, 0.827) for IVT set 2. **(b)** ROC curves for m6A (upper panel) and m5C (lower panel) for CHEUI-solo Model 1 on the IVT set 2 at different double cutoffs to separate modified and unmodified read signals. The double cutoff is indicated as an X Y pair, where detection probability > X was used to select positives and detection probability < Y was used to select negatives; all other signals being discarded. The ROC curve and double cutoffs are color matched. **(c)** The proportion of reads selected (y-axis) for each double cutoff (x-axis). **(d)** ROC AUC for CHEUI-solo model 2 and the accuracy of predicting m6A-modified (upper panel) and m5C-modified (lower panel) transcript sites calculated using independent benchmarking datasets, IVT set 2. **(e)** True positive rate (TPR) and false positive rate (FPR) for CHEUI-solo Model 2 for m6A (upper panel) and m5C (lower panel) modifications as a function of the probability cutoff (x-axis). **(f)** True Positives rates (TPR) per tool (y-axis) at different stoichiometry levels (x-axis) using an independent benchmarking dataset (IVT set 2), for m6A (upper panel) and m5C (lower panel). **(g)** False Positive Rate (FPR) (y-axis) for each tool (x-axis) returned for 512 m6A negative sites (upper panel) and 523 m5C negative sites (lower panel). Xpore had 14 false-positive site detections (FPR = 0.0273) for m681 A and 32 (FPR = 0.0611) for m5C. Epinano detected 2 false-positive sites (FPR = 0.0039) for m6A and 6 (FPR = 0.011) for m5C. CHEUI-solo had 1 false positive site detection for m5C (0.0019 FPR) and none for m6A. Nanocompore had no false positives. **(h)** Correlation between the stoichiometry predicted by each tool (y-axis) and the ground truth stoichiometry using controlled read mixtures (x-axis) for m6A (upper panel) and m5C (lower panel). We included predictions by CHEUI-solo, Xpore, Nano-RMS with the k-nearest neighbors (kNN) algorithm, and Tombo in the alternate mode (only for m5C). The Pearson correlation (r) was calculated between the predicted stoichiometries and the ground truth stoichiometry across all the sites. Other tools tested returned lower correlations and are shown in **Supp.** Fig. 5.

Inspection of the individual read probability distributions showed that modification calls with CHEUI-solo Model 1 probability close to 0.5 are more likely to be mislabeled (**Supp. Figs. 4a-4d**). We thus explored whether a double cutoff for the individual read probability would improve the accuracy. In this setting, predictions above a first probability cutoff would be considered methylated, whereas those below a second probability cutoff would be considered non-methylated, with all other read signals between these two cutoff values being discarded. Similar double-cutoff strategies have been shown before to improve the accuracy of methylation and stoichiometry estimation from DNA nanopore sequencing^38^. Amongst the configurations tested, the double cutoff 0.7 and 0.3 provided the optimal balance between accuracy gain and the number of preserved reads, with an improved area (AUC) under the receiver operating characteristic curve (ROC) for m6A (from 0.857 to 0.899) and m5C (from 0.827 to 0.877) **(Fig. 2b)**, while retaining about 73% of the reads **(Fig. 2c)**.

To train and test CHEUI-solo Model 2 for predicting the methylation probability at the transcript site level, we built *in-silico* controlled mixtures of reads, with pre-defined proportions of modified and unmodified read signals from the IVT set 1 not included previously in the training or testing of CHEUI-solo Model 1. CHEUI achieved an AUC of 0.92 for m6A and 0.953 for m5C at transcript site detection (**Fig. 2d**). Moreover, at a per-site probability > 0.99, the estimated false positive rate (FPR) on the test data was 0.00074 for m6A and 0.00034 for m5C **(Fig. 2e)**.

### CHEUI accurately detects m6A and m5C stoichiometry levels

We next compared CHEUI-solo with Nanocompore^20^, Xpore^21^ and Epinano^27^ for the ability to detect and quantify RNA modifications. To achieve this, we built positive and negative independent test datasets using read signals from IVT test 2 not used before, but with known modification state. The positive sites were built as mixtures with a pre-defined stoichiometry of 20, 40, 60, 80, and 100 percent, using 81 sites for m6A and 84 sites for m5C for each stoichiometry mixture. The negative sites consisted of 512 sites for A and 523 sites for C, using only signals from non-modified IVTs. The positive and negative sites were built by sampling reads randomly at a variable level of coverage, resulting in a lifelike coverage range of 20 to 149 reads per site. Since Nanocompore, Xpore, and Epinano required a control sample to detect modifications, a second dataset containing only unmodified signals was created for the same sites, randomly sub-sampling independent reads to the same level of coverage. We observed that the number of true positives (TP) detected by most tools increased with the site stoichiometry (**Fig. 2f**). Notably, CHEUI-solo recovered a higher number of true methylated sites compared to the other tools at all stoichiometry levels for both m6A and m5C. We next estimated the false positives by predicting with all tools on the built negative sites, using a single sample for CHEUI-solo and two independent negative samples for Xpore, Epinano, and Nanocompore. Xpore and Epinano showed the highest false positive rate (FPR) for m6A and m5C. CHEUI-solo had 1 misclassified site for m5C and none for m6A, whereas Nanocompore had no false positives (**Fig. 2g**).

We next evaluated the stoichiometry prediction in a site-wise manner. For this analysis, we included nanoRMS^28^ and Tombo^25^, which can estimate stoichiometries at pre-defined sites. Stoichiometries were calculated for the sites that were previously predicted to be modified by each tool. For NanoRMS and Tombo, the predictions for all sites were considered since these tools do not specifically predict whether a site is modified or not. CHEUI-solo outperformed all the other tools, showing a higher correlation for m6A (Pearson r = 0.839) and m5C (Pearson r = 0.839) with the ground truth (**Fig. 2h)**. CHEUI-solo was followed by Xpore (r = 0.524) and Nanocompore (r= 0.498) for m6A, and by Xpore (r = 0.556) (**Fig. 2h**) and NanoRMS (r= 0.46) (**Supp.** Fig. 5) for m5C.

### CHEUI identifies m6A modifications in cellular mRNA

We next tested CHEUI’s ability to correctly identify m6A in cellular RNA. Using DRS reads from wild-type (WT) HEK293 cells^21^ **(Supp. Table S1),** we tested 3,138,914 transcriptomic adenosine sites with a coverage of more than 20 reads in all three available replicates. Prior to any significance filtering, these sites showed a high correlation among replicates in the predicted stoichiometry and modification probability per site (**Fig. 3a)**. Analyzing the replicates together, we considered as significant those sites with prediction probability > 0.9999, which was estimated to result in an FDR nearing 0 using an empirical permutation test. After imposing this cutoff, CHEUI-solo identified 10,036 significant m6A transcriptomic sites on 3,905 transcripts, corresponding to 8,776 genomic sites **(Supp. Tables S2** and **S3**). Most of the modifications were detected on single As, with a minor proportion of AA and AAA sites predicted as modified (**Supp. Fig. 6a**). Moreover, 85.12% of the transcriptomic sites identified by CHEUI (84.5% genomic sites) had the 5′-DR**<u>A</u>**CH-3′ motif, which is a higher proportion than the 76.57% identified in m6ACE-seq and miCLIP experiments^44,45^. Interestingly, CHEUI-solo predicted m6A in 1,493 non-DRACH motifs (1,356 genomic sites), with the two most common ones being 5′-GG**<u>A</u>**CG-3′ (203 genomic sites) and 5′-GG**<u>A</u>**TT-3′ (121 genomic sites). These motifs were also the two most common non-DRACH motifs identified previously by miCLIP2 experiments in the same cell line, occurring at 245 (5′-GGACG-3′) and 96 (5′-GGATT-3′) sites^46^.

**Figure 3.**
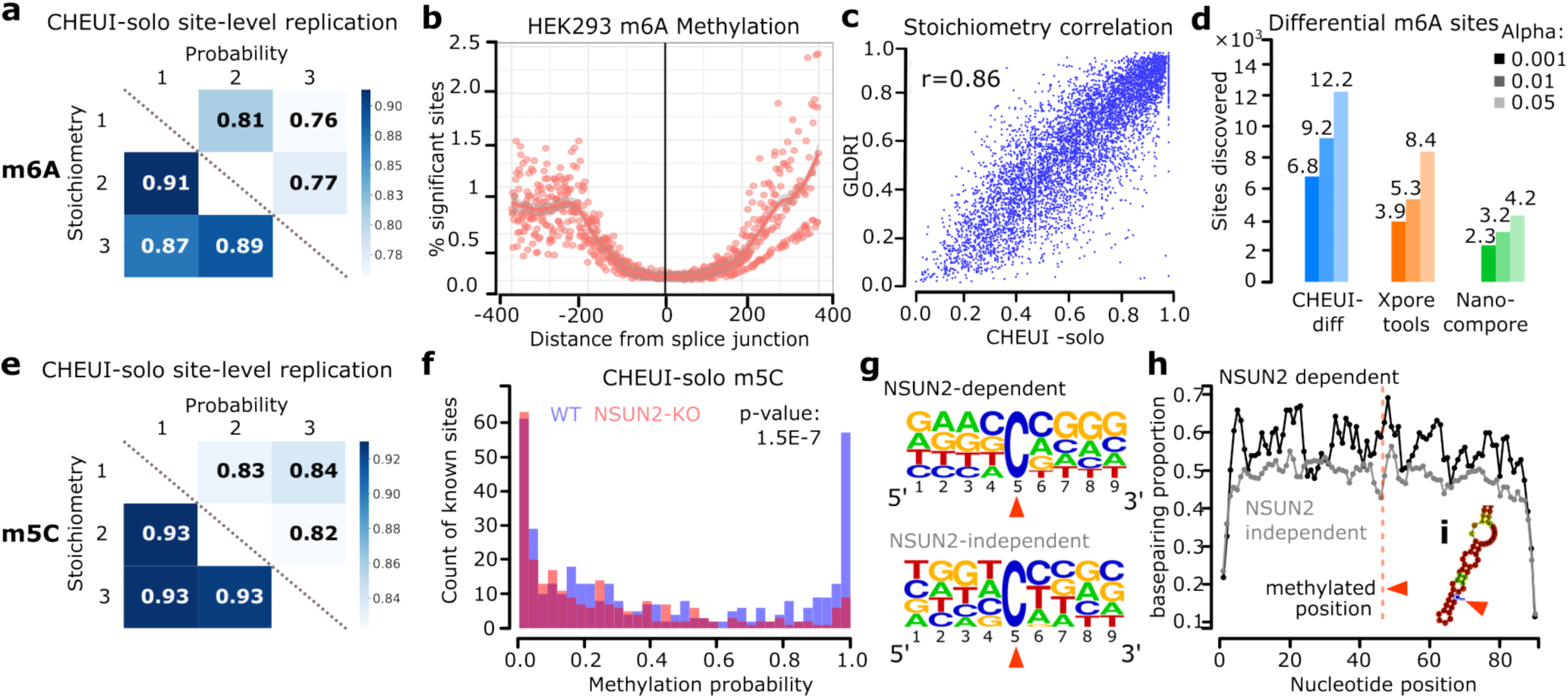
Detection of m6A and m5C in cell lines using CHEUI. **(a)** Pearson correlation values among HEK293 WT replicates for CHEUI-solo m6A stoichiometry predictions (lower diagonal) and m6A per-site probabilities (upper diagonal) for the 562,628 transcriptomic sites that had a coverage of more than 20 reads in all three replicates. **(b)** Proportion of m6A sites (predicted over tested) (y-axis) as a function of their absolute distance to the splice junction along the transcript (x-axis). **(c)** Correlation of the m6A stoichiometries in HEK293 cells estimated by the method GLORI (y-axis) and CHEUI-solo (x-axis) in 6,368 common sites. **(d)** Number of differentially modified m6A sites detected by each tool between HEK293 WT and METTL3-KO using three different levels of significance, alpha = 0.05, 0.01 and 0.001; i.e., selecting cases with adjusted p-value ≤ alpha. **(e)** Pearson correlation values among HeLa WT replicates for CHEUI-solo m5C stoichiometry predictions (lower diagonal) and m5C per-site modification probabilities (upper diagonal) for all the 497,439 tested transcriptomic sites with coverage of >20 reads in all three replicates. **(f)** Distribution of CHEUI-solo Model 2 probabilities for HeLa WT and NSUN2-KO sites also previously identified using bisulfite RNA sequencing. P-value on the upper right corner shows the result of a two-tailed Mann-Whitney U-test comparing the WT and NSUN2-KO probability distribution values **(g)** Sequence motifs for the NSUN2-dependent sites (upper panel) and the 1,000 most significant NSUN2-independent sites (lower panel) predicted by CHEUI-solo. **(h)** Proportion of base-pairing positions along 90 nucleotides centered at m5C sites predicted by CHEUI-solo. The vertical dashed red line indicates the m5C position. **(i)** Example of RNA secondary structure containing an m5C site in a stem-loop.

The m6A modification rate along mRNAs recapitulated the profile described previously, with an enrichment at the 3’ and 5’ UTRs^1,3,39^ (**Supp. Fig. 6b**). Moreover, CHEUI predictions recovered the depletion of m6A sites in the range of <200nt from the splice-sites, as was recently described^40–42^ (**Fig. 3b**). Furthermore, considering the m6A sites identified in HEK293 cells by GLORI^43^, a method based on the chemical conversion of adenosines, the 6,368 sites predicted by CHEUI and GLORI (out of the 28,865 GLORI sites with >20 nanopore reads) showed a high correlation in their estimated stoichiometries (**Fig. 3c**). We next assessed CHEUI’s false positive rate (FPR) by predicting m6A on DRS data from *in-vitro* transcribed HeLa transcriptomes^47^, which are fully non-modified. The FPR was on average 0.0003 (i.e., 3 false predictions at P>0.9999 for every 10,000 tested sites) across three replicates (**Supp. Fig. 7a**). Furthermore, CHEUI Model 2 probabilities were skewed towards zero (**Supp. Fig. 7b**) and the FPR decreased with increasing coverage (**Supp. Fig. 7c**), with most of the tested sites having coverage of 21-100 reads (**Supp. Fig. 7d**).

Next, we considered previously published DRS data from HEK293 cells with a knockout of the m6A writer METTL3 (METTL3-KO)^21^. Using CHEUI-solo predictions at individual read level, we confirmed a significant decrease in the proportion of m6A nucleotides in METTL3-KO with respect to the WT (p-value = 1.3E-254) (**Supp. Fig. 8a)**. Furthermore, using a transcriptomic site modification probability of >0.9999 as before, we corroborated the overall decrease in the proportion of modified sites along mRNAs in the KO samples (**Supp. Fig. 8b**). However, CHEUI predicted 4,603 significant m6A transcriptomic sites in METTL3-KO (**Supp. Table S4**), with 2,068 of them also present in the WT, which is consistent with recent estimates from other methods using the same cells^43,46^ and with the observation that the used METTL3-KO is not a complete allelic knockout^48^. Using an additional independent method^46^, we were able to confirm this observation (**Supp. Fig. 9**).

To compare CHEUI with other methods, we investigated the differential stoichiometry for m6A sites between HEK293 WT and METTL3-KO. CHEUI-diff showed enrichment of significant cases with higher modification stoichiometry in WT (**Supp. Fig. 8c)** (**Supp. Table S5**). In comparison with Xpore and Nanocompore, CHEUI- diff detected more sites with higher modification stoichiometry in WT at three different significance thresholds (**Fig. 3d**). CHEUI-diff also predicted a higher proportion of sites with supporting evidence from m6ACE-seq or miCLIP experiments in HEK293 cells^44,45^ (**Supp. Fig. 10a**) and containing the 5′-DR<u>A</u>CH-3′ motif (**Supp. Fig. 10b**), except at the 0.001 significance level, where 0.70 of CHEUI-diff sites and 0.71 of Xpore sites contained the motif. Comparing two METTL3-KO replicates to estimate false positives, CHEUI-diff predicted the lowest number of sites (0, 1, and 3, at the three significance thresholds, respectively) (**Supp. Fig. 10c**). In contrast, Xpore predicted over 2,000 sites at 0.001 significance and over 12,000 sites at 0.05 significance. Only 9.8% of these Xpore sites at 0.05 significance contained the 5′-DR<u>A</u>CH-3′ motif. This was a substantially lower proportion than the 46% found by Xpore in the WT *vs.* METTL3-KO comparison at the same significance level, suggesting that most of the Xpore sites in the comparison of the two METLL3-KO replicates were false positives. The overall low overlap of the nanopore-based methods with orthogonal experimental techniques suggests a different repertoire of modifications is visible to each method. This is further confirmed by the low overlap of the modification detections among diverse experimental techniques (**Supp. Fig. 11**).

### CHEUI identifies m5C modifications in cellular mRNA

We next used CHEUI to identify m5C in cell-derived RNA. To accomplish this, we used CRISPR-cas9 gene editing technology in HeLa cells to generate a knock-out (KO) of the NOP2/Sun RNA Methyltransferase 2 (NSUN2), which modifies cytosines in mRNAs and tRNAs^4,49^. The KO was confirmed by Sanger sequencing (**Supp. Fig. 12a)** and western blotting (**Supp. Fig. 12b**). The DRS **(Supp. Table S1)** yielded 2,700,022 transcriptomic sites with a coverage of more than 20 reads for the WT and 1,637,178 for the NSUN2-KO HeLa cells. Testing these sites with CHEUI-solo Model 2, prior to any significance filtering, we observed a high correlation in the predicted stoichiometry and modification probability between the replicates (**Fig. 3e**). Analyzing the three replicates together, significant transcriptomic sites were considered at probability > 0.9999, which we estimated corresponds to FDR nearing 0 using an empirical permutation test. We obtained 3,167 significant transcriptomic sites in WT (**Supp. Table S6**) and 1,841 in NSUN2-KO (**Supp. Table S7**). As above, we also assessed CHEUI’s false positive rate (FPR) using an *in-vitro* transcribed WT HeLa transcriptome^47^. We calculated an average FPR of 0.0006 (i.e., 6 false predictions for every 10,000 tested sites) across three replicates (**Supp. Fig. 7a**). As for m6A, CHEUI-solo Model 2 probabilities were skewed towards zero (**Supp. Fig. 7b**) and the FPR decreased with increasing coverage (**Supp.** Fig. 7c), with most of the tested sites having coverage of 21-100 reads (**Supp. Fig. 7d**).

As we observed before for m6A, the prediction of two or more adjacent m5C sites was rare, and most of the predictions were individual m5C sites (**Supp. Fig. 13a**). We then compared CHEUI-solo m5C calls with a union set of 7,918 sites previously detected in HeLa using bisulfite RNA sequencing (bsRNA-seq) in three independent studies^4,8,49^ (**Supp. Fig. 13b**). From these sites, 372 (4.7%) had >20 nanopore reads and could be tested by CHEUI. These sites showed a higher probability than sites without bsRNA-seq evidence (**Supp. Fig. 13c**). Additionally, CHEUI-solo detection probabilities on this union set of 372 sites were significantly higher in WT compared with NSUN2-KO (**Fig. 3f**). Further validating this result, a permutation analysis to compare the probability of these sites against the background distribution of probabilities in the same samples confirmed that CHEUI-solo returned higher probability modification detection values in the WT samples than expected by chance (p-value = 0.001) (**Supp. Fig. 13d**). In contrast, the NSUN2-KO did not show this enrichment (p-value = 0.025) (**Supp. Fig. 13e)**. In contrast to what we found for m6A, looking at individual nucleotides with CHEUI-solo Model 1 we observed only a mild reduction in the proportion of m5C over the total cytosine occurrences in the NSUN2-KO compared with the WT (p-value = 1.2E-35) (**Supp. Fig. 14a**). Moreover, the profile of significant m5C sites along mRNAs did not change between the WT and NSUN2-KO (**Supp. Fig. 14b**). These results are consistent with reports showing that a fraction of m5C sites in mRNA are NSUN2-independent^4,49^, which have been proposed to be regulated by NSUN6^50,51^.

To investigate NSUN2-dependent sites, we used CHEUI-diff to identify differentially modified sites between WT and NSUN2-KO (**Supp. Table S8**). This yielded 186 potential NSUN2-dependent unique genomic sites, 18 of which were previously identified by bsRNA-seq. In contrast, Nanocompore and Xpore only found 4 and 11 overlaps with bsRNA-seq sites in this comparison, respectively, while they predicted many more sites transcriptome-wide (**Supp. Fig. 15**). Furthermore, the 186 potential NSUN2-dependent sites showed similarity to the previously described sequence motif for NSUN2-dependent sites: 5′-m5CNGGG-3′^49^ (**Fig. 3g**). We also identified 1,250 NSUN2-independent sites, defined as those predicted in WT but with no significant change relative to KO, which showed a different motif (**Fig. 3g**). Encouragingly, these NSUN2-independent sites occurred in genes significantly enriched in mitotic cell cycle function (p-value = 6.206E-5) and processes (p-value = 1.594E-4), which agrees with previous findings for genes with NSUN2-independent sites^52^.

To further assess the validity of our predictions, we investigated the likelihood of RNA secondary structure formation in their vicinity. Consistent with previous studies^4,49^, canonical base-pair probabilities were higher in NSUN2-dependent sites compared to NSUN2-independent sites (**Fig. 3h and 3i**), and the potential base-pairing arrangements suggested a higher occurrence of stem-loops at around 5 nt downstream of the NSUN2-dependent m5C site (**Supp. Figs. 16** and **17).** Further supporting CHEUI results, NSUN2-dependent sites identified previously by bsRNA-seq^49^ showed significantly higher stoichiometry differences between WT and NSUN2-KO than all other m5C sites (**Supp. Fig. 18)**.

To further investigate the correspondence between nanopore-based predictions and bsRNA-seq, we performed DRS and bsRNA-seq on the RNA obtained from newly designed IVT templates s generated to be either fully m5C modified or non-modified (**Supp. Table S9**). Applying permissive parameters to the analysis of bsRNA-seq data (Methods), we estimated a conversion rate of 0.9987 in the non-modified samples and 0.0269 in the modified samples. However, from the 4,423 C sites on the IVT templates, 99.55% were covered in the non-modified sample but only 72.12% in the modified sample. In contrast, >99% of the C sites were covered with more than 20 nanopore reads in both samples, and hence visible to CHEUI (**Supp. Table S9**). Unlike for the cellular transcriptome, we cannot use the permutation analysis to select a CHEUI probability cutoff. We thus calculated at various probability cutoffs the recall, as the proportion of m5C-sites detected in the modified IVTs, and the false positive rate, as the proportion of predicted m5C sites in the non-modified IVTs. At P>0.99, the FPR was less than 1%, with a recall of 65.22% (**Supp. Table S9**). Moreover, at P>0.99, CHEUI predicts 824 of the 1,219 sites missed by bsRNA-seq, with 26 of these 9-mers also predicted by CHEUI but not by bsRNA-seq in the HeLa WT data. These 26 9-mers included sites with 2 and 3 Cs together. This suggests that at C-rich sites, nanopore sequencing may present an advantage over bsRNA-seq in the identification of m5C.

### Impact of other modifications on the prediction of m6A and m5C

To test if other modifications could impact the detection of m6A or m5C in individual read signals, we tested CHEUI on the signals from IVTs containing other modifications not used for training, namely, 1-methyladenosine (m1A), hydroxymethylcytidine (hm5C), 5-formylcytidine (f5C), 7-methylguanosine (m7G), pseudouridine (Y) and inosine (I)^26^. All read signals were processed for each 9-mer centered at A or C as before, with the modification either at the same central base (m1A and m6A for A, and m5C, 5fC, and hm5C for C) or in the neighboring bases in the 9-mer (Y, m7G, I, m1A, m6A for C; or Y, m7G, I, m5C, 5fC, hm5C for A) (**Fig. 4a)**. As a general trend, the proportion of signals containing other modifications predicted as positives by CHEUI recapitulated the results for signals without any additional modifications (**Fig. 4b**). This was the case for all modifications, except for predictions by the m6A model in signals containing m1A, a chemical isomer of m6A, which followed a similar trend as m6A (**Fig. 4b**, upper panel).

**Figure 4.**
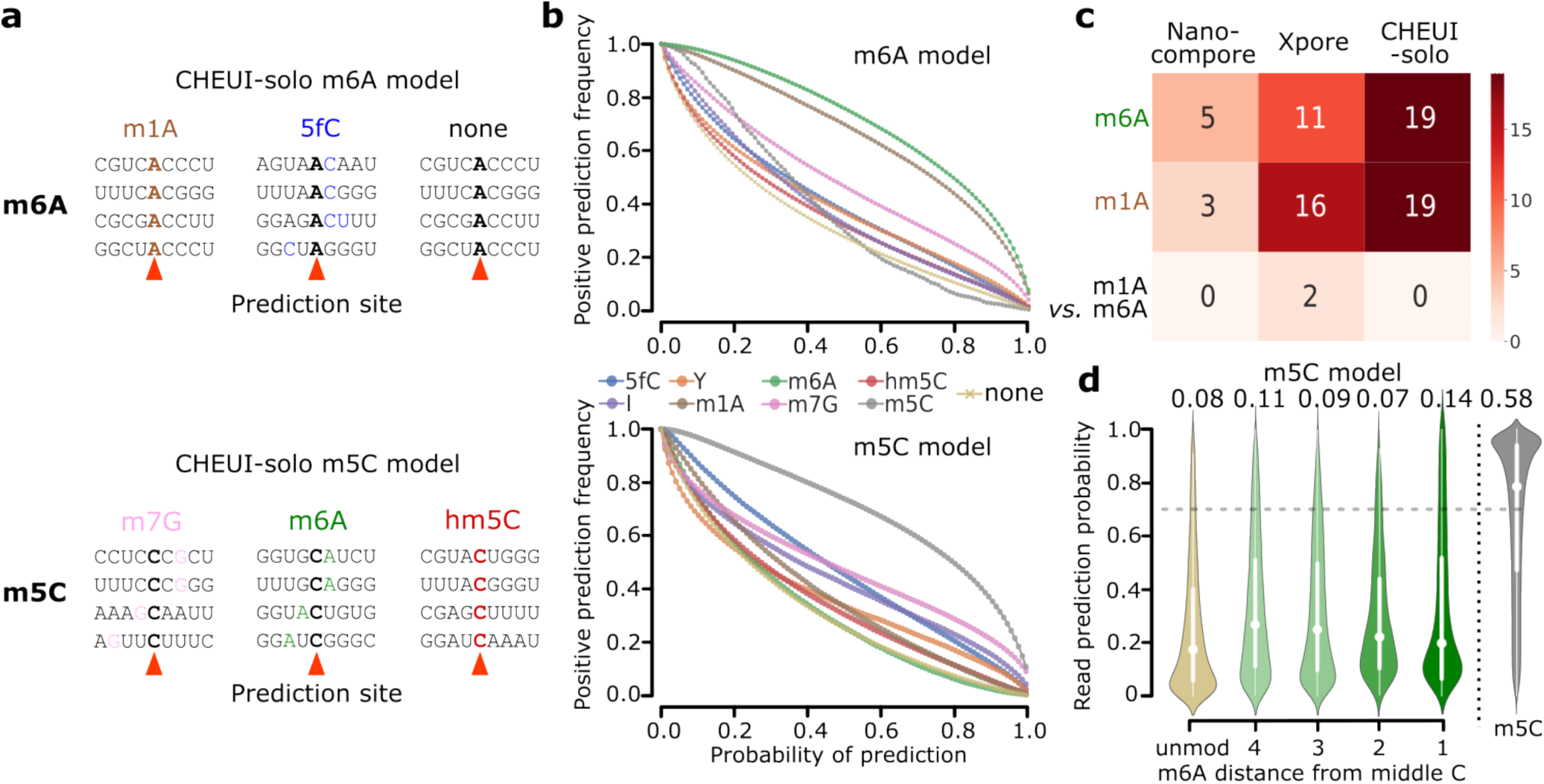
Impact of nearby presence of other RNA modifications on the detection accuracy of m6A and m5C in nanopore signals. **(a)** Examples of some configurations for which CHEUI-solo Model 1 was tested in individual reads for m6A (upper panel) and m5C (lower panel) using signals from IVTs containing other modifications. **(b)** The number of read signals identified as m6A- (upper panel) and m5C- (lower panel) containing by CHEUI-solo Model 1 (y-axis) at different values of the probability cutoff (x-axis). **(c)** The number of significant sites identified by each tool (x-axis) in each of the conditions (y-axis). The ‘m6A’ and ‘m1A’ row show the number of sites with 100% stoichiometry predicted as m6A by each method. For Nanocompore and Xpore, these were calculated by comparing each sample against the unmodified sample. The ‘m6A *vs.* m1A’ row shows the number of sites with a significant difference between the two modified samples. For CHEUI, the number of sites was calculated as those detected only in one of the samples. **(d)** CHEUI-solo’s detection probability of m5C at individual read level (y-axis) using IVT set 2 read signals at 9-mers with a single C at the center and considering various configurations: 9-mers with no m5C (unmod), 9-mers with m6A present at relative position 1, 2, 3, or 4 from the central C, and 9-mers with a modified middle C (m5C). The proportion of read signals identified as modified with probability > 0.7 is indicated above each distribution.

To investigate whether m1A misclassification might be due to the similarity between the m1A and m6A nanopore signals, we used Xpore and Nanocompore to test the discrimination of m6A and m1A without any *a priori* assumption about the modification type. We used 81 9-mers centered at A and made all possible pairwise comparisons among three sets of read signals: one with no modifications, one with all signals having m1A, and one with all signals having m6A. Coverage per site ranged between 21 and 324 reads, with a median coverage of 62 reads. When comparing m6A or m1A against unmodified signals, Xpore identified significant differences for 11 and 16 sites, Nanocompore detected 5 and 3 sites, and CHEUI m6A model predicted 19 sites in both cases (**Fig 4c**). In the comparison of m6A against m1A read signals, Xpore found a significant difference in only two of the sites, whereas Nanocompore found none (**Fig. 4c**). These results suggest that the DRS signals for these two isomers may be indistinguishable with the current statistical models and/or pore chemistry (**Supp. Fig. 19**). To further address the m6A and m1A DRS signal similarity, we retrained the CHEUI-solo m6A model using m1A signals as negatives and m6A signals as positives. Although this new model achieved accuracy comparable to the original one in the separation of m6A from non-modified signals (**Supp. Fig. 20a**), it showed a trade-off between accurately detecting m6A and correctly separating m6A from m1A (**Supp. Fig. 20b**), further indicating existing limitations to separate these isomeric RNA modifications using the nanopore signals.

We further assessed how the presence of one modification may impact the detection of the other at short distances in individual reads. We analyzed the detection of m5C at 9-mers in non-modified individual reads and in reads where m6A was present nearby, using reads from the IVT test 2 datasets. The proportion of false positives (0.07-0.14) for the CHEUI m5C Model 1 when m6A was 1-4 nt away was similar to the background proportion with no modification (0.08). The proportion of false positives for m6A detection in the presence of a nearby m5C modification (0.08-0.12) was also similar to the background level (0.13) (**Supp. Fig. 21**).

### Transcriptome-wide analysis suggests m6A and m5C co-occurrence in individual mRNA molecules

We next used CHEUI’s ability to identify m6A and m5C from the same sample to investigate the potential co-occurrence of modifications in a mammalian transcriptome. Using WT HEK293 data, we calculated whether individual reads covering two predicted modified transcriptomic sites presented specific modification combinations (i.e., m6A-m5C, m6A-m6A, m5C-m5C) more frequently or at a similar rate in comparison with random pairs of modifications sites from different transcripts. Intriguingly, we observed that read-level modification co-occurrence, defined as the proportion of molecules with both sites having the same modification status, was higher than expected by chance for m6A and m5C modifications (**Fig. 5a**). This increased co-occurrence was observed at short distances (<5 nt) as well as long distances (>5 nt). Given the observed partial influence of nearby modifications in the prediction of m5C and m6C described above, we decided to perform an additional test. At each position, we compared the observed co-occurrence with the expected value calculated by independently permuting the modification status across the reads. As a result, m5C upstream of m6A (i.e., 5’-m5C…m6A-3’) showed a significant co-occurrence at short distances (5-8 nt) and at longer distances (11-12 nt) (**Fig. 5b**), whereas m6A upstream of m5C (i.e. 5’-m6A…m5C-3) showed a significant co-occurrence only at longer distances (13-15 nt) (**Fig. 5b**). We also found that m6A-m5C sites with 40-60% stoichiometry showed the most significant overrepresentation compared to the values expected by chance (**Supp. Fig. 22**). The co-occurrence of m6A-m6A or m5C-m5C was also higher than expected at short distances (1-4 nt) but was close to expected values at longer distances (5-15 nt) (**Supp. Fig. 23**). Furthermore, discarding m6A and m5C sites at distances <5 nt from each other, we also observed an enrichment of transcripts harboring both modifications relative to the total number of m6A and m5C transcriptomic sites, both in HEK293 (**Fig. 5c**) and HeLa (**Supp. Fig. 24**).

**Figure 5.**
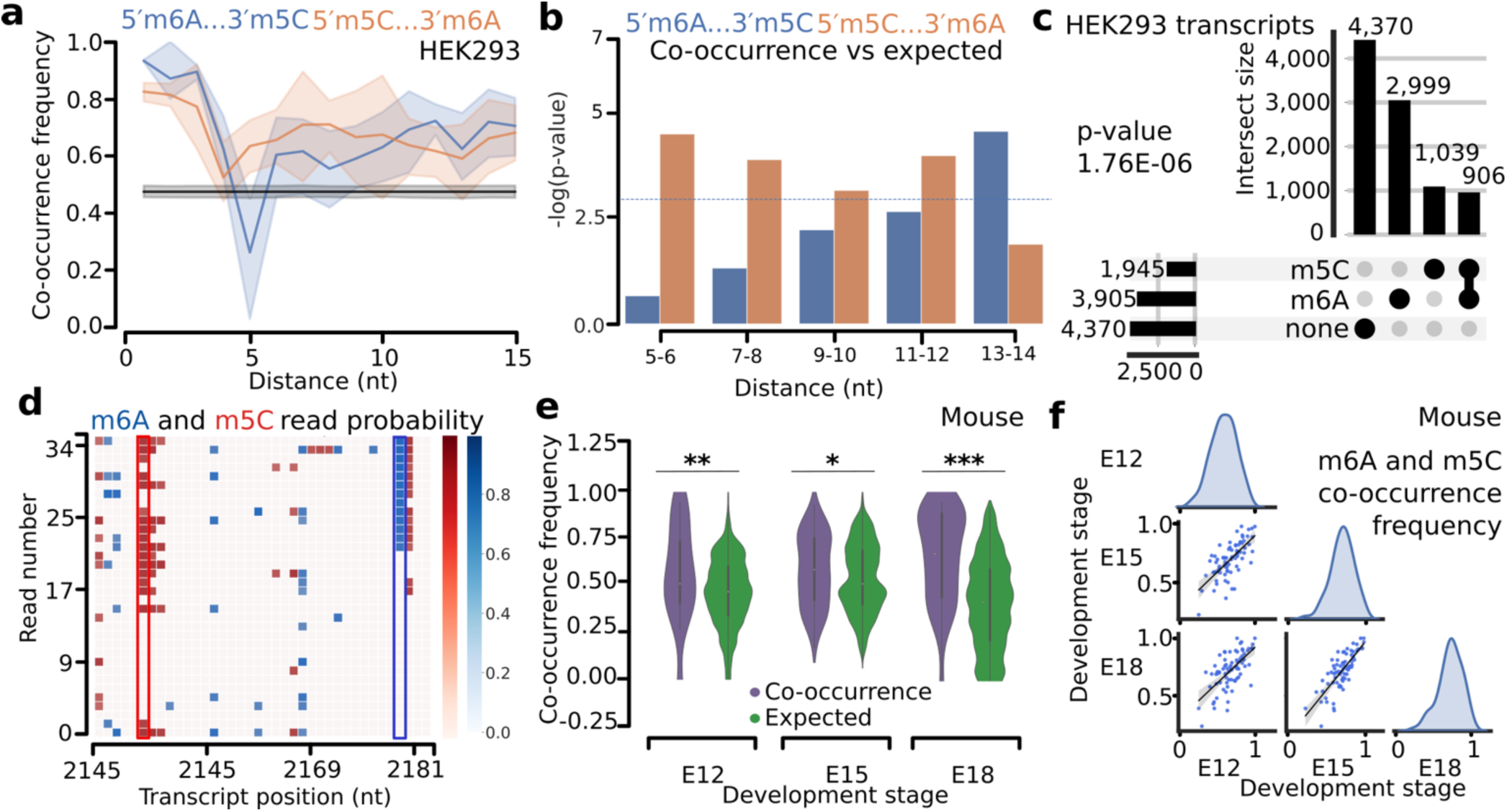
Coordinated occurrence of m6A and m5C in RNA *in vivo*. **(a)** Co-occurrence (y-axis) of m6A and m5C modifications at the read level at various relative distances (x-axis). Co-occurrence is calculated by counting the number of reads where two sites have the exact same modification state, either both modified (CHEUI-solo Model 1 probability > 0.7) or both unmodified (CHEUI-solo Model 1 probability < 0.3) divided by the total number of reads covering both sites. Pairs of sites with A upstream of C are depicted in blue and pairs of sites with A downstream of C are shown in red. Dark blue and orange lines indicated mean co-occurrence values, and the shades indicate the 95% confidence intervals of the mean at each distance. The black line and grey shades indicate the mean and 95% confidence interval values of the random co-occurrence values. Distances are measured as the difference between positions of the two modified nucleotides, e.g., 5′-<u>m6A</u>NNNN<u>m5C</u>-3′ are at the relative distance of 5 nt. **(b)** Statistical significance (y-axis) of the comparison between the observed and expected co-occurrences at different distances for the pairs of m6A and m5C sites (x-axis). Co-occurrences and expected values in the x-axis were considered in groups of two consecutive distances. Observed and expected co-occurrences were compared using a one-sided Mann-Whitney U-test. A horizontal blue line indicates p-value = 0.05. The expected co-occurrences were calculated by permuting the modification status of reads in each site independently. **(c)** Number of human protein-coding transcripts containing m6A and m5C sites, only one of the modifications, or none, in HEK293. Only cases with m6A and m5C at a distance of 5 nt or more were considered. The p-value corresponds to a two-tailed Fisher’s test for an increased observed co-occurrence. **(d)** Region of the transcript ENST00000258214 (gene *CCDC102A*) showing m6A and m5C modifications occurring in individual reads (y-axis) at various transcript positions (x-axis). The blue scale shows CHEUI’s detection probability at the read level for m6A and the red scale depicts the same but for m5C. The positions identified as modified by CHEUI-solo Model 2 are 2,150 nt for m5C with 0.66 stoichiometry, and 2,179 nt for m6A with 0.72 stoichiometry (highlighted). 26 molecules exhibited m5C, whereas 21 molecules exhibited m6A, while 18 molecules harbored both sites. The co-occurrence of 0.78 is calculated using the molecules that harbor both sites. **(e)** Distribution of the observed and expected read-level co-occurrences of m6A and m5C (in any relative order) (y-axis) in the RNA of mouse embryonic cortex at three development stages, E12, E15, and E18 (x-axis). Both orientations (m6A-m5C and m5C-m6A) were considered together in the range of relative distances of 5-15 nt. One-sided Mann-Whitney U-tests comparing the observed and expected distributions resulted in p-values of 0.0044, 0.0148, and 1.5586E-14, respectively for the E12, E15, and E18 developmental stages. Expected random distributions were obtained by permuting the individual read methylation states at each site independently. **(f)** Correlations between the co-occurrence values at the individual-read level for pairs of m6A and m5C sites in a pairwise comparison between RNA from each of the mouse frontal cortex developmental stages, E12, E15, and E18. Pearson correlation between E12 and E15 was r=0.68 (p-value 7.8E-12), between E12 and E18 was r=0.64 (p-value 4.2E-10), and between E15 and E18 was r=0.78 (p-value 1.3E-16). Density distributions of the co-occurrence values are additionally shown.

To examine how CHEUI resolves m6A and m5C co-occurrences in RNA molecules, we visualized a region of 38 nt from 34 RNA molecules derived from the transcript ENST00000258214 of the gene *CCDC102A,* which codes for a protein component of the myosin complex (**Fig. 5d**). These RNA sequences present high confidence predictions by CHEUI-solo Model 2 (probability > 0.9999) for m6A (position 2,179 nt of the transcript) with 0.72 stoichiometry and m5C (position 2,150 nt of the transcript) with 0.66 stoichiometry, with 78% of the individual molecules containing both modifications (**Fig. 5d**). While nucleotides adjacent to these modified sites had a high modification probability at the level of individual reads by CHEUI-solo Model 1 (probability > 0.7), the corresponding transcript reference sites were not considered significant by CHEUI-solo Model 2. Generally, consecutive modified sites were rarely detected using our defined cutoff for CHEUI-solo Model 2 (probability >0.9999) (**Supp. Figs. 6a** and **13a**).

An intriguing question is the possibility of a coordinated m6A and m5C occurrence in a physiological context, where RNA modifications play an important role. We decided to study m6A and m5C during brain development, where m6A has been reported to be relevant^53^. We collected cortex tissue from wild-type mice at three different embryonic stages E12, E15 and E18, and performed DRS of 3′ poly(A)^+^ RNA (**Supp. Fig. 25**) (**Supp. Table S1**). We tested a total of 1.4M to 2.2M transcriptomic ‘A’ sites and 1.2M to 2M transcriptomic ‘C’ sites. Using the probability cutoff of > 0.9999, we obtained 2,876 to 6,040 m6A sites and 1,390 to 2,180 m5C sites (**Supp. Tables S2** and **S10**), with modification rates along mRNAs similar to those observed for the cell lines (**Supp. Fig. 26**). We found that in all three conditions, m6A and m5C modifications at distances of 5 nt or more co-occurred in transcripts significantly more often than expected by the random incidence of the two modifications (**Supp. Fig. 27**).

The pairs of methylated sites (m6A-m5C and m5C-m6A) in each condition showed a wide variation in co-occurrence at the level of individual reads, but the global co-occurrence values were significantly higher than expected by chance at each of the three developmental stages (**Fig. 5e**). Moreover, read-level co-occurrences were higher than expected by chance at distances 5-15 nt and at low and intermediate stoichiometries (**Supp. Fig. 28**). Furthermore, co-occurrence values of m6A-m5C sites showed a high correlation among the three embryonic stages, suggesting that the co-occurrence of modifications is transcript-specific and conserved across this developmental timeline **(Fig. 5f)**. The conservation of the co-occurrence was apparent even for the sites of low stoichiometry across developmental points, which can be exemplified by a 35 nt region from the transcript ENSMUST00000014438 (gene *Ndufa2*), where co-occurring m6A and m5C sites were found 13 nt apart (**Supp. Fig. 29**). While the modification frequency in these sites was moderate at about 30%, the m6A-m5C and m5C-m6A co-occurrence in molecules were 0.961, 0.957 and 0.913 for the E12, E15 and E18, respectively, consistent with the identified high conservation across conditions.

## DISCUSSION

We have developed CHEUI for the transcriptome-wide identification of m6A and m5C from the same sample, both in individual molecules as well as in transcriptome reference sites, together with their stoichiometry quantification, without requiring a KO/KD or an otherwise control sample. To assess the expected performance of CHEUI, we performed an in-depth benchmarking using *in-vitro* transcribed RNA for which we knew the methylation status in each read. This was particularly effective for the assessment of stoichiometry, which is challenging to test in cellular RNA where a complete knowledge of the modification status of all RNA molecules is generally not available. Using controlled mixtures of modified and non-modified reads, we tested variable coverage and stoichiometry values using different nanopore-based approaches. These analyses showed that CHEUI accomplishes high sensitivity, precision, and modification stoichiometry accuracy compared to other nanopore-based tools. We further used the IVT strategy to compare side-by-side bisulfite RNA-sequencing (bsRNA-seq) and CHEUI for the identification of m5C on a ground truth dataset. This showed that CHEUI presents a trade-off between the recall of m5C in modified IVTs and the false positive rate in non-modified IVTs. This analysis also indicated a potential limitation of bsRNA-seq for the detection of m5C in C-rich contexts, which can be recovered using nanopore sequencing.

For the analysis of cell transcriptomes, given the large number of sites tested, we used very strict cut-offs to maintain a low expected false discovery rate (FDR). While these strict cutoffs resulted in a reduction of CHEUI sensitivity, many of sites identified by orthogonal methods had high CHEUI probabilities that were below the set thresholds. Relaxing these thresholds would recover more sites found by orthogonal techniques but at the cost of introducing potential false positives. This limitation may stem from the variability of the nanopore signals, where the differences between modified and non-modified reads are often comparable to the differences observed in the population of non-modified reads. Improvements in the prediction models and in the RNA sequencing chemistry can potentially facilitate the identification of m6A and m5C at higher sensitivity.

In cell transcriptomes, CHEUI and other tested nanopore-based methods showed a low correspondence with orthogonal experimental methods. For m6A, we observed a low overlap with CLIP-based sites, which also showed a low overlap among different experiments in the same cell models. We also observed a low overlap with bsRNA-seq for m5C. Furthermore, CHEUI and other nanopore-based methods generally detected many potential m5C sites there were not present in the bsRNA-seq datasets. These results suggest that there are biases and differences in detection rates associated with each technology and that much is yet to be learned about the full distribution of modifications in mRNA. Independent validation experiments of the modified sites detected only from nanopore reads will be necessary to confirm these predictions or establish whether they are due to other sources of nanopore signal variation. Further strategies to address these discrepancies between technologies could involve identifying consensus approaches that combine multiple experimental sources or nanopore-based methods trained on a wider range of experimental inputs.

We observed that CHEUI and other nanopore-based methods tested could not accurately separate the positional isomers m1A and m6A. Visual inspection of the signals for m6A and m1A in the same k-mer contexts suggests that they deviate in the same way from the signals corresponding to unmodified nucleotides. In contrast, m5C and hm5C, which have different chemical groups attached to the same position, may be visually distinguished from each other and from the unmodified nucleotides. Difficulties to separate the isomeric m1A and m6A have also been described with other technologies^54^. More sophisticated predictive models including additional features to the nanopore signal, such as neighbouring sequence motifs or secondary structure, could overcome the observed limitation.

CHEUI’s capacity to predict two modifications in the same sample enabled us to measure the co-occurrence of m6A and m5C in transcripts. While our systematic analysis of signals shows that at distances of >5 nt the co-occurrence of m6A and m5C can be reliably identified in the reference transcriptome and in individual molecules, we observed a residual mutual signal interference of the modifications at distances less than 5 nt. This effect on the individual read prediction is exacerbated in ribosomal RNA molecules since they are highly and densely modified. This limitation could be addressed by incorporating additional predictive features or developing new models trained on datasets with defined combinations of adjacent modifications.

The mechanisms underlying the identified co-occurrence of modifications in reads and across transcripts remain to be elucidated. A possibility could be a crosstalk between RNA modification enzymes, whereby the binding of RNA by readers or writers for one modification may drive the deposition or removal of the other. However, a preliminary analysis of 9-mers devoid of As yielded the same m5C modification rate in WT and METTL3-KO cells. Similarly, the m6A modification rate in 9-mers devoid of C’s was the same in WT and NSUN2-KO cells (data not shown). While these results may not show the full picture given the existence of other NSUN enzymes, they suggest that if there was a link between the m6A and m5C deposition, this might not occur across the transcriptome. Further experiments would be required to settle this question.

There are other reasonable explanations for non-random modification co-occurrence that do not require the interaction of the methylation machineries. Co-occurring modifications at a single-molecule level may represent the relics from the ‘history’ of the RNA molecule, which acquired the modifications by passing through certain processing steps or points of cellular response^58^. Such epitranscriptomic relics may contain entangled modifications of various types, combinations of which can be characteristic of a subpopulation of the cell’s RNA with a shared history. Another possibility is an enhanced accessibility of the RNA to the methylation enzymes induced by one or the other modification, possibly in contrast to the cases where such accessibility is not present due to the mRNA localization or translation. A more evolutionary-inspired possibility is the correction of function, whereby once a modification is introduced, it enhances or compensates for the functional effects of a pre-existing modification. Using more targeted experiments with methodologies that can identify more than one modification from the same sample, such as CHEUI, can potentially provide further insights into the co-occurrence of modifications in individual molecules and open new opportunities to study the functions embodied by the epitranscriptome.

## METHODS

### Nanopore signal preprocessing

The nanopore sequencing data was preprocessed using the following steps prior to running CHEUI. First, the FAST5 files were basecalled using Guppy. IVT datasets were basecalled with Guppy version 4.0.14. Data from mouse (E12, E15, E18) and cell lines (WT and METTL3-KO in HEK293 cells, and WT and NSUN2-KO in HeLa) were basecalled using Guppy version 5.0. Reads were then aligned to the corresponding reference transcriptome using Minimap2^59^. The genome and annotation references used were GRCh38 and Gencode v38 for the human data, and GRCm39 and Ensembl v104 for the mouse data. For the IVT reads, options ‘-ax map-ont -k 5’ were used, whereas for human and mouse transcriptomes, the options ‘-ax map-ont –k14’ were used. Reads were then filtered to select the best match for each read using samtools *-F 2324*^60^. Nanopolish’s (version 0.13.2)^17^ *eventalign* was then used to align the read signals to the matched transcript references using the options ‘--scale-events -- signal-index --samples --print-read-names’. Nanopolish *eventalign* output consists of 5-mers along the transcript reference and a list of signal values for each of those 5-mers. Although each 5-mer is given in the 5′ to 3′ orientation, the list of signals per 5-mer is ordered in the 3′ to 5′ orientation. To process the signals in the right 5′ to 3′ orientation, we thus flipped the signals per 5-mer before concatenating the signals from overlapping 5-mers. All the (per read) signals for every 5 overlapping consecutive 5-mers, together with the read ID and sequence, were then used to create the input for CHEUI-solo Model 1.

### CHEUI-solo Model 1

#### Model description

CHEUI-solo Model 1 is a convolutional neural network (CNN) modified from the Jasper model^61^. The model architecture (**Supp. Fig. 3**) was implemented using Keras^62^ and Tensorflow^63^. The input for this CNN is defined as follows. For a given position of interest, e.g., adenosine (A) for the m6A model or cytosine (C) for the m5C model, given the 9-mer centered at that position, i.e., NNNN(A|C)NNNN, CHEUI uses the signals corresponding to the five consecutive overlapping 5-mers including that middle position of interest. The number of signals is, in general, variable and was fixed before being used as input for the CNN model. The signals for each 5-mer are then converted into a 20-length vector by dividing the values into 20 segments preserving their order and calculating the median value for each segment. If a 5-mer contained more than 20 values, the values were divided into 20 equal subsets, and the median value of each subset was used. If the event had fewer than 20 values, the median was appended to these values until it reached 20 values. As a result, each 9-mer was then mapped to a vector of 5×20 = 100 signal values, which is used as input for CHEUI-solo Model 1. CHEUI also uses as input the distance between the observed and the expected signal for every 5-mer. The expected signal is built using the k-mer model from Nanopolish^17^, which describes the signal value for each 5-mer in the absence of modifications. For each of the 5 overlapping 5-mers in the observed signals, each expected value was repeated 20 times to obtain a vector of expected values of length 100. Then, a vector of length 100 with the absolute distances between the components of the expected and the observed signal vectors is calculated. These vectors of observed signals and absolute distances are used as input for CHEUI-solo Model 1. Of note, CHEUI-solo Model 1 does not use the actual k-mer (k=9) sequence, only the vector of observed signals and the vector of distances, providing a high level of abstraction form the sequence context.

#### Training and testing of CHEUI-solo Model 1

CHEUI-solo Model 1 was trained using read signals generated from *in-vitro* transcript (IVT) data^26,27^ to produce one model for each modification, m6A or m5C. The positive training set contained m6A (or m5C) in place of the canonical nucleotides, i.e., every A was replaced by m6A (or every C by m5C)^27^. For both models, the ‘negative’ sets were made from read signals from IVTs but with no modifications. For both modifications, we constructed non-overlapping datasets for training (IVT train 1), validation (IVT validation 1), and testing (IVT test 1, IVT test 2) (**Supp. Table S11**). Datasets IVT train 1, IVT test 1, IVT validation 1 were built from publicly available reads^27^, using non-overlapping signal reads for each dataset that could share the same 9-mer sequence contexts. IVT train 1 was composed of 9-mers with any number of As (or Cs) in the modified and unmodified sequences. IVT validation 1, used for parameter optimization, was composed of 9-mers containing only one A (or C) at the center of the 9-mer. IVT test 1, which was used to test sensor generalization, was also composed of 9-mers with only one A (or C) at the center. On the other hand, IVT test 2, used to test k-mer generalization, was built from independent IVT experiments^26^. IVT test 2 contained non-overlapping signal reads and included 9-mer contexts that were not present in the other train, test, or validation datasets. IVT test 2 was also composed of 9-mers with only one A (or C) at the center of the 9-mer. Importantly, the training and testing was performed on individual read signals.

Binary cross-entropy was used as the objective function, AMSGrad was used as the optimizer, and the Nvidia Tesla V100 was used to accelerate computing. Training was performed for 10 epochs and for every 200,000 read signals the accuracy, precision, recall and binary cross-entropy loss were calculated on the IVT validation 1 set along with the parameters of the model at that stage. After 10 epochs, there was no improvement on the validation accuracy, so the training was terminated. Accuracy was defined as the proportion of correct cases, i.e. (TN+TP)/(TN+TP+FN+FP); precision was calculated as the proportion of predicted modifications that were correct, i.e. TP/(TP+FP) and recall as the proportion of actual modifications that were correctly predicted, i.e., TP/(TP+FN); where TP = true positive, FP = false positive, TN = true negative, FN = false negative. Binary cross-entropy was defined as

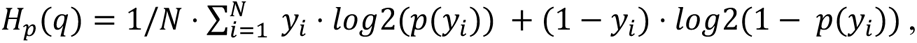

where *y_i_* = 1 for a modified base in a specific position of a read and 0 otherwise, and *p(y_i_)* is the posterior probability from the Model 1.

### CHEUI-solo Model 2

#### Model description

CHEUI-solo Model 2 is a binary classifier implemented as a CNN like for Model 1. CHEUI-solo Model 2 takes as input the distribution of probabilities generated by Model 1 for all read signals at a given transcriptomic site, i.e., a position in a reference transcript, and predicts the stoichiometry and probability of that site being methylated (m6A or m5C). Model 2 assumes that the distribution of the individual-read probabilities at a given transcriptomic site originates from two classes, one with a subset or all reads having high Model 1 probabilities (modified site), and a second one with low Model 1 probabilities (unmodified site).

#### Model training and testing

CHEUI-solo Model 2 was trained using controlled mixtures of modified and unmodified reads not used previously for training, validation, or testing of CHEUI-solo Model 1. These controlled mixtures were built to comprise a wide range of values for coverage and stoichiometry, and with a high proportion of low coverage and low stoichiometry sites, to mimic what was previously observed in transcriptomes^4,49,64^. The new read signals were processed as described before and used to make predictions with CHEUI-solo Model 1. The training set for Model 2 consisted of mixtures of modified and unmodified reads from IVTs^27^ with their corresponding Model 1 probabilities. To model the low stoichiometry and coverage values, the training sites were built as follows: 1) a site was chosen to be modified or unmodified with 50% probability; 2) if unmodified, a coverage was chosen randomly between 0 and 100, using a linear decay, i.e., the higher the coverage, the less likely it was to be selected, and the per-read probabilities were assigned at random from the pool of unmodified signals; 3) if, on the contrary, the site was selected to be modified, the coverage and stoichiometry of the site were chosen using the same linear decay as before, with high coverage and stoichiometry values less likely to be chosen. The linear decay was implemented using the *random.choices* function from the general python distribution using the weights *(10 - coverage) × 0.01 + 0.9* as argument. Weights indicate the relative likelihood of each element on the list to be chosen, with each incremental unit of coverage or stoichiometry corresponding to a decrease in their weight by one unit. Using this procedure, we generated approximately 1.5M synthetic sites per modification with variable coverage and stoichiometry. These sites were randomly split into training and testing in a 9:1 proportion.

### Comparison with other tools

#### Tools selected for comparison

We chose tools available for each specific benchmarking comparison. We used Epinano^27^, which implements a linear regression with two samples, one depleted of modifications to detect outliers, i.e., observations with large residuals, to identify modifications. We used *EpiNano-Error,* which combines all types of read errors (mismatches, insertions and deletions) in pairwise mode. We also used NanoRMS^28^, which does not predict modified sites but uses predictions from another method to calculate the stoichiometry with a sample comparison approach. Specifically, NanoRMS uses the signals processed by Tombo or Nanopolish and implements a supervised *k-*NN method based on the sample labels, or an unsupervised method based on *k*-means with *k*=2, to separate modified and unmodified signals. For NanoRMS, the stoichiometry was calculated from the proportion of reads from the WT sample in the modified cluster, divided by the total number of WT reads. We also tested Nanocompore^20^, which uses the assignment of raw signals to a transcriptome reference with Nanopolish and the mean current value and mean dwell time of all signals per 5-mer, and then compares the distributions for all read signals aligning on the same site between two conditions. Nanocompore then fits a Gaussian mixture model with two components to the data and performs a statistical test to determine whether each cluster is significantly associated with a sample. We also tested Xpore^21^, which operates similarly to Nanocompore, using the assignment of raw signals to the transcript reference with Nanopolish and comparing the mean current values between two or more conditions for each transcriptomic site. Xpore uses information from unmodified k-mers as a prior for Gaussian distributions and variational Bayesian inference to infer the mean and variance of each distribution. After fitting the data into clusters, Xpore labels clusters with values closer to the expected unmodified signals as unmodified and then performs a statistical test on the differential modification rates between samples and assigns a p-value per site. We also tested Tombo in *sample comparison* mode, which performs a statistical test comparing the signal values between two conditions; and Tombo in *alternative mode,* which predicts a proportion of m5C modification per transcriptomic (not individual read) site, although it does not provide a score or a probability for the modification calls.

#### Controlled IVT mixtures for benchmarking

To create a controlled and independent dataset to benchmark the accuracy in the prediction of stoichiometry and transcript-site modification, we used the reads from Jenjaroenpun, P. *et al.*^26^ not used in the previous tests to generate mock ‘WT’ and ‘KO’ samples. The mock ‘WT’ sample was generated by randomly sampling reads from the modified and unmodified sets to create multiple stoichiometry mixtures with 20, 40, 60, 80, and 100 percent.

The mock ‘KO’ sample was created by randomly sampling reads from the unmodified pool of reads. We ran Epinano, Nanocompore, Xpore, and CHEUI, using default parameters to predict RNA modifications. Epinano, Nanocompore, and Xpore were run using the generated WT and KO mock samples. CHEUI was run using only the generated WT sample, as it does not require a KO/KD or control sample. Predicted sites were considered at three levels of significance or alpha values, i.e., predicted sites were considered significant if, after correcting for multiple testing, the adjusted p-values were ≤ alpha, where alpha = 0.05, 0.01, 0.001.

Transcript-site predictions, i.e., the methylation state of a position in the reference sequence, in the IVT-based mixtures were classified as positive if they had a probability > 0.99 from CHEUI-solo Model 2, and negative otherwise. Nanocompore, Xpore, Epinano, and CHEUI were run using thresholds recommended by the documentation for each tool. For Xpore, sites containing a k-mer (k=9) centered in adenosine, in the evaluation of m6A, or a cytosine, in the evaluation of m5C, that had a predicted p-value lower than 0.05 were considered significant. For Nanocompore, the same selection of k-mers centered in adenosine or cytosines was done, and sites with a p-value lower than 0.05 were selected as positives. For Epinano, we used Guppy version 3.0.3 and *EpiNano-Error* with the combined errors *Epinano_sumErr* method to detect modifications, as recommended in the Epinano documentation. We then used the linear regression model and ‘unm’ or ‘mod’ from the ‘linear model residuals z score prediction’ column to classify sites as unmodified or modified, respectively.

To estimate the false positive rate for Epinano, Nanocompore, and Xpore we evaluated the number of sites each tool predicted as modified when comparing two sets of reads with no modifications. For CHEUI, we used only one of those datasets with no modifications. We evaluated all sites with A or C, regardless of whether they had other As or Cs nearby in the same k-mer (k=9) sequence context. In contrast, to determine the true positive rate and stoichiometry, we only evaluated k-mers (k=9) containing one centered m6A and no additional As, or one centered m5C and no additional Cs to avoid the influence of having two or more modified nucleotides affecting the tested site, since the IVTs were built with all nucleotides of one type either modified or not modified.

#### Stoichiometry benchmarking

Stoichiometries were calculated in the following way. Given a modified site identified by CHEUI-solo Model 2 at an annotated transcript position in a given sample, the stoichiometry is calculated as the proportion of reads covering that site that have the site identified as modified according to CHEUI-solo Model 1. For the analyses presented, we used the probability by CHEUI-solo Model 1 > 0.7 to tag a site as modified at the individual read level, and < 0.3 to tag the site as unmodified at the individual read level, discarding calls with probability values in the range [0.3, 0.7]. Stoichiometry was only calculated in transcriptomic sites predicted as positively modified by CHEUI, i.e., with a CHEUI-solo Model 2 probability of > 0.9999. For Xpore, we used the values of the column ‘mod_rate_WT-rep1’, which we interpreted as the modification rate of the mock ‘WT’ sample. In the case of Nanocompore, we used the column ‘cluster_counts’ that contains the number of WT and KO reads that belong to the two clusters, one modified and the other unmodified. Stoichiometry was then calculated as the percentage of modified reads in the ‘WT’ sample, i.e., we divided the number of WT reads in the modified cluster by the total number of WT reads. We also included NanoRMS with *k*-NN and *k*-means for the stoichiometry comparison. In this case, since NanoRMS only predicts the stoichiometry on sites predicted by another method and since Epinano predicted very few sites in our test set, we applied NanoRMS to all tested sites (81 for m6A and 84 for m5C) to obtain a more unbiased assessment. The percentage of modified reads per site was obtained from the NanoRMS output tables, dividing the number of modified reads in the WT by the total number of WT reads. Finally, Tombo assesses every site and gives a fraction of modified reads but does not specify the site as modified or not. As most of the sites had a fraction of modified reads above 0, even for the unmodified sample (75 out of 84 sites), we only considered Tombo for the stoichiometry comparisons.

### Testing m6A and m5C accuracy in read signals with other modifications

For this test, we used the Nanopore signals for the IVT transcripts from Jenjaroenpun, P. *et al.*^26^. Each dataset contained either unmodified signals, or signals for modified nucleotides with m6A, m5C, 1-methyladenosine (m1A), hydroxy-methylcytidine (hm5C), 5-formylcytidine (5fC), 7-methylguanosine (m7G), pseudouridine (Y), and Inosine (I) modifications. We considered all 9-mers centered at A or C in the IVT reads containing modifications other than m6A (for A-centered 9-mers) or m5C (for C-centered 9-mers). Thus, the modifications were either at the same central base (m1A and m6A for A; and m5C, 5fC, and hm5C for C) or in neighboring bases (Y, m7G, I, m1A, m6A for C; or Y, m7G, I, m5C, 5fC, hm5C for A). We used CHEUI-solo Model 1 to predict m6A in the middle A or m5C in the middle C for all these read signals, to determine the influence of these other modifications on CHEUI’s ability to correctly separate A from m6A and C from m5C.

### CHEUI-solo for transcriptome-wide analyses

Reads from the three replicates for each condition WT HeLa, NSUN2-KO HeLa, WT HEK293, and METTL3-KO HEK293 were aligned to the Gencode v38 transcriptome (GRCh38) using minimap2 as described above. CHEUI-solo (Model 1 and Model 2) was run on pooled replicates from each condition, except when comparing replicates within the same condition. In each case, CHEUI-solo Model 1 was run on all the reads, whereas CHEUI-solo Model 2 was run only on transcriptomic sites with the coverage of more than 20 reads. This produced a methylation probability and estimated stoichiometry in all tested transcriptomic sites. To establish a probability cutoff of significance for CHEUI-solo Model 2, we calculated the probability distribution of modified sites expected by chance, without a biological signal. To do so, in each given condition, we shuffled all read signals across all transcriptomic sites, maintaining the same number of transcriptomic sites and the same coverage at each site. We then run CHEUI-solo Model 2 over these sites with the new read signal distributions obtained after shuffling the reads. For each tested probability cutoff, the proportion of candidate transcriptomic sites selected as methylated from the shuffled configuration was considered as an estimate of the false discovery rate (FDR). Using this approach, we found that a probability cutoff of 0.9999 for CHEUI-solo Model 2 would yield an FDR = 0 for m6A, and an FDR = 0.000384 for m5C. We thus consider modified transcriptomic sites the ones having a Model 2 probability equal to or higher than 0.9999 for both modifications. CHEUI-solo (Model 1 and 2) was also applied as above to three replicates of DRS of in-vitro transcribed WT HeLa cells. Predicted m6A or m5C sites with Model 2 probability > 0.9999 were considered as false positives.

### Comparison with other methods for m6A detection in HEK293 cell lines

Xpore, Nanocompore and CHEUI-diff were used to call differential RNA modifications on all A sites, using 3 WT and 3 KO replicates for HEK293. CHEUI-diff was run on sites that had >20 reads in both conditions, WT and KO. We used three distinct levels of significance: 0.05, 0.01, and 0.001. For Xpore and CHEUI-diff, FDR correction was performed with Benjamini-Hochberg procedure. Since Nanocompore already provides adjusted p-values, the threshold was applied without FDR correction. To compare the transcriptomic sites identified as m6A in WT, we selected those sites predicted by each method to have increased stoichiometry in the WT. By default, CHEUI-diff does not test sites where the difference in stoichiometry between the two conditions is less than 0.1 in its absolute value. For Xpore, we used the module *xpore postprocessing* to filter the output. To calculate the potential number of m6A false positives we used each tool to compare two replicates from the same KO condition with the highest number of reads, METTL-KO replicates 2 and 3. The KO was used instead of the WT samples to minimize the chances of including variably modified m6A sites that may occur in WT samples. To compare the nanopore-based predictions with m6A transcriptomic sites with previous evidence we employed the union of m6ACE-seq and miCLIP sites^44,45^.

### CHEUI application to the signals derived from RNA of NSUN2-KO and WT HeLa cells

CHEUI-solo (Models 1 and 2) was run by pooling together three replicate samples from each cell line, WT and NSUN2-KO HeLa. Information about previously identified m5C sites in HeLa was collected from three different bisulfite RNA sequencing (bsRNA-seq) experiments^4,8,49^ and the union of these three sets was considered for subsequent comparisons. The probabilities of the modification calls derived from CHEUI-solo Model 2 corresponding to sites with orthogonal evidence were compared between WT and NSUN2-KO using a two-tailed Mann-Whitney U-test.

The permutation analysis to test the enrichment of high probability calls in the candidate sites detected by bsRNA-seq was performed in the following way. First, we calculated how many bsRNA-seq candidate sites were tested by CHEUI-solo (total sites) and how many of these were the ‘high probability sites’, defined as those having Model 2 probability of >0.99. Then, we randomly sampled the same number of transcriptomic sites tested with CHEUI-solo Model 2 and counted how many of these were high-probability sites. We repeated this procedure 1,000 times and calculated an empirical p-value.

Sequence logos were computed using WebLogo (https://weblogo.berkeley.edu/logo.cgi). To study the propensity of secondary structure formation around NSUN2-dependent and -independent m5C sites, we used RNAfold 2.4.18^65^. We estimated base-pairing probabilities in the region covering 90 nucleotides centered over the m5C site (45 nt on either side). For each sequence, we calculated the nucleotide positions that had pairwise interactions with other nucleotides according to RNAfold. At each position, we then calculated the proportion of nucleotides with interactions with respect to the total number of sequences. These proportions were plotted separately for the WT and NSUN2-KO samples. The enrichment of functions and processes associated with genes with modifications was assessed using g:Profiler^66^.

### Bisulfite RNA sequencing analysis

We performed RNA bisulfite treatment (bsRNA-seq) following the protocol from Johnson et al.^67^. There were no deviations done for the protocol except the fact that a GE50 spin column was used for the removal of the excess of bisulfite reagent (sodium bisulfide and hydroquinone) instead of a GE25. This protocol was applied to RNA fully modified with m5C or non-modified obtained from in-vitro transcripts (IVTs) m1, m2, m3, m4 (Supp. Table S9) built from four non-overlapping fragments from the mouse canonical pre-rRNA (∼13kb long). Sequencing of the bisulfite-treated samples was performed with Illumina. For the analysis of the Illumina reads from the bisulfite-treated data we used meRanTK^68^, adjusting the parameters to make it more permissive to m5C detection: the edit distance was changed from the default 2 to 200, the number of Cs per Illumina read was changed from the default 3 to 200, the minimum methylation ratio of a single C needed for methylation was changed from the default 0.2 to 0, and the minimum coverage at a given reference site above which methylation call is performed was changed from the default 20 to 0. The same modified and non-modified RNA samples were used to perform nanopore DRS.

### CRISPR-Cas9 knockout (KO) of NSUN2 in HeLa cells

#### HeLa cell lines and culture

HeLa cells (human cervical cancer) were obtained from ATCC (cat. no CCL-2) and confirmed *via* short tandem repeat (STR) profiling with CellBank Australia. Cells were grown in DMEM medium (Gibco) supplemented with 10% FBS and 1× antibiotic-antimycotic solution (Sigma) and passaged when 70–90% confluent. HeLa cell cultures were tested to be negative for mycoplasma contamination prior to their processing for gene editing.

#### Guide sequence design

Two CRISPR (cr)RNAs were designed, targeting the 5′-proximal (exon 2 crRNA “AGGCUACCCCGAGAUCGUCA”) and 3′-proximal (exon 19 crRNA “AAUGAGAGUGCAGCCAGCAC”) regions of the gene. Gene sequences from Ensembl (Asia server) were processed *via* CCTop^69^ to check for efficacy and predict potential off-target cleavage effects. The two sequences with highest predicted efficacy and minimal off-target effects were selected as crRNA and ordered as Alt-R CRISPR-Cas9 crRNA from Integrated DNA Technologies (IDT).

#### Ribonuclear protein preparation

2.5 µM of NSUN2 exon 2 crRNA was combined with equimolar amounts of NSUN2 exon 19 crRNA and annealed with 5 µM Alt-R CRISPR-Cas9 trans-activating CRISPR (tracr)RNA, ATTO 550 (IDT) in 10 µl of 1× IDT Duplex Buffer. The ribonuclear protein (RNP) assembly reaction was then performed by combining 0.575 µM of the annealed crRNA:tracrRNA with 30.5 pmol of IDT Alt-R S.p. Cas9 Nuclease V3 in 2.2 µl Neon Transfection System ‘R’ resuspension buffer (Invitrogen) for 5 minutes at 37 °C; the resultant mixture was kept at room temperature until transfection.

#### Transfection

Electroporation was conducted using Neon Transfection System (Invitrogen) and following the manufacturer’s protocol, with the following modifications. HeLa cells were resuspended in Neon Transfection System ‘R’ resuspension buffer (Invitrogen) to a concentration of 2.8×10^7^ per ml. For each electroporation reaction, 2×10^5^ cells prepared as above were incubated with 1× v/v RNP at 37 °C for 5 minutes, before being electroporated at 1,005 volts, 35 milliseconds, with 2 pulses. Two reactions were seeded per well of a 24-well plate. Cells were recovered in complete medium under standard incubation conditions of 37 °C and 5% v/v CO_2_ for 24 to 36 hours.

#### Single cell sorting

Cells were sorted for singlets and ATTO 550 positivity on a FACSAria II Cell Sorter (BD) hosted at the Flow Cytometry Facility of the John Curtin School of Medical Research, the Australian National University. Although all singlets were positive when compared with negative controls, only cells with high intensity ATTO 550 (>10^33^ RFU) were sorted into 96-well plates for subsequent culturing. Cells were maintained in complete media and expanded to 6-well plates for genomic DNA (gDNA) extraction upon reaching 70% confluency.

#### Amplicon analysis

The gDNA was extracted by incubating cell pellets with 30 µl of in-house rapid lysis buffer (40 µg Proteinase K, 10 mM Tris-HCl pH 8.0, 1 mM EDTA, 0.1% v/v Tween-20) at 56 °C for 1 hour followed by denaturation at 95 °C for 10 minutes. Amplification of NSUN2 gene was conducted with standard protocols under 35 cycles in Mastercycler Nexus (Eppendorf), using Q5 High-Fidelity DNA Polymerase (New England BioLabs) and 5 µl of extracted gDNA. Amplicons were purified with ExoSAP-IT (Applied Biosystems) and sequenced on an AB 3730xl DNA Analyzer, by the ACRF Biomolecular Resource Facility (BRF) from the John Curtin School of Medical Research, Australian National University, following the manufacturer’s protocol (Applied Biosystems 2002). Sequencing data was analyzed manually using SnapGene software (from Insightful Science; available at https://www.snapgene.com/) to confirm alteration of the target loci.

#### Protein analysis

Cells were grown in DMEM medium (Gibco) supplemented with 10% FBS and 1× antibiotic-antimycotic solution (Sigma) and passaged when 70-100% confluent. Unmodified wild-type (WT) and NSUN2 KO cells were scraped in 200-500 μl of protein extraction buffer (50 mM Tris pH 7.5 at 25 °C, 5 mM EDTA, 150 mM NaCl, 21.5 mM MgCl_2_, 10% glycerol, 1% v/v Triton X-100, 1× Complete EDTA-free Protease Inhibitor Cocktail (Sigma)) and incubated for 10 minutes on ice, then incubated for 30 minutes at 4°C on a rotator. The mixture was centrifuged at 13,000 g for 10 minutes at 4 °C. The supernatant was transferred to a clean tube, and used immediately, or stored at –80 °C. Total protein concentration was then estimated by taking a Qubit measurement via Protein Assay Kit (Thermo Fisher Scientific) following the manufacturer’s instructions. 30 μg of total protein was loaded on NuPage 4-12% w/v Bis-Tris Protein Gels (Invitrogen), and proteins were electrophoretically separated using NuPAGE MES SDS Running Buffer under conditions recommended by the manufacturer. Separated proteins were transferred onto PVDF membrane using iBlot 2 Transfer Stacks, PVDF, mini (Thermo Fisher Scientific, cat. no. IB24002), following manufacturers’ instructions. The membrane was blocked in Odyssey Blocking Buffer (LI-COR, cat. no. 927-40000) and probed with primary antibodies: anti-NSUN2 (1:1,000; Proteintech, cat. no. 20854-1-AP), anti-ACTB (1:1,000; SantaCruz, cat. no. sc-47778 AF790). The membranes were then incubated with the anti-rabbit-IR-Dye680 secondary antibody (1:10,000; LI-COR, cat. no. 925-68071) and scanned using the Odyssey CLx Imaging System (LI-COR). The KO’s effect was assessed by the specific intensity alteration of the fluorescent signal of the respective band with mobility corresponding to that expected of NSUN2.

#### Extraction of polyadenylated RNA from HeLa cells

Three each ⌀10 cm plates with WT and NSUN2-KO HeLa cells at 80% confluency were washed twice in ice-cold PBS and scraped in 500 µl of denaturing lysis and binding buffer (100 mM Tris-HCl pH 7.4, 1 % w/v lithium dodecyl sulfate (LiDS), 0.8 M lithium chloride, 40 mM EDTA and 8 mM DTT; LBB). The cell lysate was thoroughly pipetted with 200 µl tip until the sample viscosity was reduced, and pipetting was seamless. 500 µl of oligo(dT)_25_ magnetic beads (New England Biolabs) suspension was then used per replicate. The beads were washed with 1 ml of LBB twice, each time collecting the beads on a magnet and completely removing the supernatant. Upon washing, the oligo(dT)_25_ beads were resuspended in the cell lysate and placed in a rotator set for 20 rpm at 25 °C for 5 minutes, followed by the same rotation at 4 °C for 30 minutes. The suspension was briefly spun down at 12,000 g, separated on a magnet, and the supernatant was discarded. The beads were then resuspended with 1 ml of wash buffer (20 mM Tris-HCl pH 7.4, 0.2 % v/v Titron X-100, 0.4 M lithium chloride, 10 mM EDTA and 8 mM DTT; WB) and washed on a rotator set for 20 rpm at 4 °C for 5 minutes, using 3 rounds of washing. The beads were collected on a magnetic rack and the supernatant was discarded. The wash procedure was repeated three times. The elution was carried out stepwise. Washed bead pellet was first resuspended in 50 µl of the elution buffer (25 mM HEPES-KOH, 0.1 mM EDTA; HE). The suspension was heated at 60 °C for 5 minutes to facilitate the elution, and the eluate was collected upon placing the bead-sample mixture on a magnetic rack, separating the beads, and recovering the clean supernatant. The resultant pellet was next resuspended in another 50 µl of HE buffer, and the process was repeated.

The eluates from oligo(dT) bead extraction were combined and further purified using AMPure XP SPRI beads (Beckman Coulter Life Sciences) generally according to the manufacturer’s recommendations. Briefly, the eluate samples were supplemented with 1.2× volumes of the SPRI bead suspension in its standard (supplied) binding buffer, and the resultant mixture incubated at room temperature for 5 minutes with periodic mixing. The SPRI beads were brought down by a brief 2,000 g spin and separated from the solution on a magnetic rack. The supernatant was removed, and the beads were resuspended in 1 ml of 80 % v/v ethanol, 20 % v/v deionized water mixture and further washed by tube flipping. The bead and solution separation procedure were repeated. The ethanol washing process was repeated one more time. Any remaining liquid was brought down by a brief spin and removed using a pipette, and the beads were allowed to air-dry while in the magnetic rack for 2 minutes. The purified RNA was then eluted in deionized water and the RNA content was assessed using absorbance readout *via* Nanodrop and fluorescence-based detection *via* Qbit RNA high sensitivity (HS) assay kit (Thermo Fisher Scientific). RNA was then stored frozen at –80 °C until downstream processes were required.

#### RNA DRS Library Preparation for HeLa samples

The library preparation generally followed the manufacturer’s recommendations. 650-800 ng of RNA from HeLa cells were used for each 2× library preparation within every replicate (with all recommended volumes doubled-up) with direct RNA sequencing kit (SQK-RNA002) as supplied by Oxford Nanopore Technology. The modifications were that Superscript IV RNA Polymerase (Thermo Fisher Scientific) was used, RNA Control Standard (RCS) was omitted, and RNasin Plus (Promega) was included at 1 U/ µl in all reaction solutions until the SPRI purification step after the reverse transcription reaction. The final adaptor-ligated sample was eluted in 40 µl.

### Embryonic mouse brain development experiments

#### Brain tissue extraction

Mice (strain C57BL/6J) were dissected on embryonic day (E) E12, E15 and E18. All procedures were conducted in accordance with the Australian National University Animal Experimentation Ethics Committee (protocol number A2019/46). Pregnant females were cervically dislocated, and (male and female) embryos were extracted in cold sterile PBS. The frontal area of the cortex, i.e., the pallium, was then dissected with micro-knifes under a Zeiss STEMI 508 stereomicroscope and tissue samples were immediately placed in a 1.5 ml microcentrifuge tube (Eppendorf) containing 300 µl of denaturing lysis and binding buffer (100 mM Tris-HCl pH 7.4 at 25 °C, 1 % w/v lithium dodecyl sulfate (LDS), 0.8 M lithium chloride, 40 mM EDTA and 8 mM DTT; LBB). Samples were immediately agitated by vigorous pipetting until near-complete tissue dissolution, flash-frozen on dry ice and stored at – 80 °C until downstream processes were required.

#### Polyadenylated RNA extraction from the denatured brain development samples

About 150 mg of the original (wet weight without denaturing buffer) of the cortex tissue was used per extraction. Upon defrosting, the tissue/LBB mixture was thoroughly pipetted with 200 µl tip until the sample viscosity was reduced, and pipetting was seamless. 500 µl of oligo(dT)_25_ magnetic beads (New England Biolabs) suspension was used per replicate. The beads were washed with 1 ml of LBB twice, each time collecting the beads on a magnet and completely removing the supernatant. Upon washing, the oligo(dT)_25_ beads were resuspended in the tissue/LBB mixture and placed in a rotator set for 20 rpm at 25 ^°^C for 5 minutes, followed by the same rotation at 4 °C for 30 minutes. The suspension was briefly spun down at 12,000 g, separated on a magnet, and the supernatant was discarded. The beads were then resuspended with 1 ml wash buffer (20 mM Tris-HCl pH 7.4, 0.2 % v/v Titron X-100, 0.4 M lithium chloride, 10 mM EDTA and 8 mM DTT; WB) and washed on a rotator set for 20 rpm at 4 °C for 5 minutes, 3 wash rounds in total were performed. For each wash, the beads were collected on a magnetic rack and the supernatant was discarded. The elution was carried out stepwise. Washed bead pellet was first resuspended in 50 µl of the elution buffer (25 mM HEPES-KOH, 0.1 mM EDTA; HE). The first suspension was heated at 60 °C for 5 minutes to facilitate the elution, and the eluate was collected upon placing the bead-sample mixture on a magnetic rack, separating the beads, and recovering the clean supernatant. The resultant pellet was next resuspended in another 50 µl of HE buffer, and the process was repeated. The eluates were then combined and subjected to an additional solid-phase reversible immobilization (SPRI) bead purification step as described in the ‘Extraction of polyadenylated mRNA from HeLa cells’ sub-section above and stored frozen at –80 °C until downstream processes were required.

### MinION flow cell priming and DRS

Nanopore sequencing was conducted on an Oxford Nanopore MinION Mk1B using R9.4.1 flow cells for 24-72 hours per run, depending on the flowcell exhaustion rate. Tthe flow cells were left at 25 °C for 30 minutes to reach ambient temperature. The flow cells were then inserted into the MinION Mk1B and a quality check was performed to ensure that the pore count was above manufacturer warranty level (800 pores). Prior to the sample loading, the priming solution (Flush Buffer mixed with Flush Tether) was degassed in a vacuum chamber for 5 minutes. A similar approach was repeated when loading the RNA library. The run set up on the loaded libraries was performed according to the recommended running options using MinKNOW software (Version 4.3.25). The SQK-RNA002 sequencing option was selected, and the bulk file output was switched from OFF to ON to export the complete data. For real-time assessment of the quality of the run, the output FAST5 files were base called in-line with sequencing using the MinKNOW-provided Guppy software.

### RNA abundance analysis of the embryonic mouse brain tissue development sequencing data

Basecalled reads were aligned to the mouse reference genome (GRCm39) using minimap2 v2.1.0 (parameters: ‘minimap2 -ax splice -k14 -B3 -O3,10 --junc-bonus 1 --junc-bed’). During alignment, splice junction coordinates were provided to minimap2 in BED format using the ‘junc-bed’ flag to improve the accuracy of the spliced alignments. Splice junction BED files were generated using minimap2 paftools.js gff2bed function, using the the gene structure reference (Ensembl 2014 mouse GTF). Primary genomic alignments were assigned to genes using Subread featureCounts v2.0.1 in stranded, long-read mode (using parameters --primary -L -T 48 -s 1 --extraAttributes ‘gene_biotype, gene_name’). DESeq2 v1.26.0^71^ was used to obtain log-normalized gene counts. PCA plots were generated from regularized log transformed gene counts, using DESeq2’s plotPCA function.

### Liftover of transcriptomic to genomic sites and calculation of metatranscript coordinates

We used our R2Dtool^72^ (https://github.com/comprna/R2Dtool) to perform positional annotation of the CHEUI Model 2 RNA methylation calls, and to transpose the methylation predictions from transcriptomic to genomic coordinates. First, the R2Dtool script *cheui_to_bed.sh* was run with default parameters to convert the CHEUI methylation calls to a bed-like format (i.e., tab-delimited, where column 1 represents the reference sequence, column 2 represents interval start, and column 3 represents interval end). Next, the R2Dtool *R2_annotate* command was used with default parameters and the relevant GTF annotation (Ensembl v104 / GRCm39 GTF for mouse and Gencode v38 / GRCh38 for human) to perform positional annotation of the bed-like CHEUI Model 2 methylation calls. Positional annotation included metatranscript coordinates, and the distances from a given site to the nearest upstream and downstream splice junctions annotated (if applicable) in the same transcript where the modified site was predicted. Finally, the R2Dtool *R2_lift* command was run with default parameters to transpose the annotated methylation calls from transcriptomic coordinates (i.e., position on a specific transcript) to genomic coordinates (i.e., position on a specific chromosome).

### RNA methylation metatranscript plots

The absolute distance (in nucleotides) and relative metagene position (as a fraction of the overall UTR or CDS length) of each methylation site with respect to the reference transcript isoform were calculated using R2Dtool^72^. The relative meta-transcript coordinates were derived as previously described^73^, placing the modifications along three equal-sized segments of length L. Position 0 represents the transcript start site (TSS), position L represents the CDS start, position 2L represents the CDS end, and position 3L represents the polyadenylation site (PAS). For our graphical representation, we used L=40. Meta-transcript plots showing the abundance of tested and significant sites, alongside the proportion of significant sites per tested region, were made using ggplot2 (https://ggplot2.tidyverse.org/).

### Co-occurrence of modifications in transcripts and reads

To study the co-occurrence of modifications in annotated transcripts, we considered all protein-coding transcripts (‘mRNAs’) with at least two tested sites, i.e., by default having 20 or more reads at each site. For the co-occurrence of m6A and m5C, we partitioned these mRNA transcripts into four sets according to whether they contained two significant m6A and m5C sites, only one of the modifications, or had no significant sites (even though both were tested). Based on this partition, we performed a two-tailed Fisher’s exact test to determine whether the association of m6A and m5C in transcripts was higher than expected. To study the co-occurrence of modifications in reads, we considered those transcripts with two modified sites at a relative distance from 1 to 15 nt. We then calculated the co-occurrence as the proportion of reads with both modifications, i.e., the number of reads that at both sites have the same modification status divided by the total number of reads considered. To calculate the expected level of co-occurrence in the same sample, we calculated the co-occurrence for 1,000 pairs of modified sites located in different transcripts. For this analysis, we discarded any possible reads and sites of the ribosomal RNAs (rRNAs) (only present in the mouse data). It is known that rRNAs are hypermodified in multiple positions. Considering our analysis of the effects of other modifications on the identification of m6A and m5C, we expect these to be affected by the other modifications.

## Data availability

The synthetic sequence templates from Liu et al.^27^ were obtained from the NCBI Gene Expression Omnibus (GEO) database under the accession number GSE124309. The nanopore read signals for the in-vitro transcribed (IVT) RNAs obtained from these synthetic sequence templates with m6A, m5C, or no modifications, were obtained from NCBI Sequence Read Archive (SRA) under accessions PRJNA511582 and PRJNA563591. Nanopore data for the synthetic transcripts from Jenjaroepun et al.^26^ was obtained from The Sequence Read Archive (SRA) accession SRP166020. Nanopore data for HEK293 WT and METTL3-KO samples from Pratanwanich et al.^21^ was obtained from the European Nucleotide Archive (ENA) under accession PRJEB40872. Data from the m6ACE-seq experiments from Koh et al.^45^ was obtained from the NCBI Gene Expression Omnibus (GEO) under accession number GSE124509. Nanopore data for HeLa WT and HeLa NSUN2 KO and for the embryonic mouse brain tissues produced in this work have been deposited at NCBI GEO under accession GSE211762. Nanopore sequencing and bisulfite RNA sequencing data for the IVT RNAs is available at NCBI GEO under accession GSE253150.

## Code availability

CHEUI is freely available from https://github.com/comprna/CHEUI under an Academic Public License

R2Dtool (v1): https://github.com/comprna/R2Dtool

Nanocompore (v1.0.0rc3-2): https://github.com/tleonardi/nanocompore

Xpore (v0.5.4): https://github.com/GoekeLab/xpore

Epinano (v0.1-2020-04-04): https://github.com/novoalab/EpiNano

Tombo (v1.5): https://github.com/nanoporetech/tombo

NanoRMS (Downloaded on the 2^nd^ of July 2021): https://github.com/novoalab/nanoRMS

Keras (v1.1.2): https://github.com/keras-team/keras

Tensorflow (v2.4.1): https://github.com/tensorflow

Minimap2 (v2.1.0): https://github.com/lh3/minimap2

Nanopolish (v0.13.2): https://github.com/jts/nanopolish

RNAfold (v2.4.18): https://www.tbi.univie.ac.at/RNA/

## Supplementary Tables description

Supp. Table S1: Description of the samples used in this study.

Supp. Table S2: Number of m6A and m5C CHEUI sites in each of the samples from Supp. Table S1.

Supp. Table S3: Significant CHEUI-solo m6A and m5C sites in HEK293 WT.

Supp. Table S4: Significant CHEUI-solo m6A sites in HEK293 METTL3-KO.

Supp. Table S5: CHEUI-diff significant differential m6A sites between WT and KO.

Supp. Table S6: Significant CHEUI-solo m6A and m5C sites in HeLa WT.

Supp. Table S7: Significant CHEUI-solo m5C sites in HeLa NSUN2-KO.

Supp. Table S8: CHEUI-diff significant differential m5C sites between WT and KO.

Supp. Table S9: Comparison of CHEUI with bsRNA-seq on the same RNA samples.

Supp. Table S10: Significant CHEUI-solo m6A and m5C sites in mouse embryonic brain at three development stages.

Supp. Table S11: Number of IVT inputs used for training and testing of CHEUI.

## Acknowledgements

We are grateful to the personnel from the Biomolecular Resource Facility at JCSMR (ANU), and particularly to Tiffany Cripps, Lachlan Morrison, Carolina Correa Ospina and Stephanie Palmer, for their assistance with DNA validation sequencing. We are also grateful to the personnel of the Ecogenomics and Bioinformatics Lab, a joint initiative of the Research School of Biology (ANU) and Commonwealth Scientific and Industrial Research Organisation, and particularly to Niccy Aitken and Ashley Jones for their continued support and feedback regarding ONT sequencing.

