## Supplementary Figures for "Prediction of m6A and m5C at single-molecule resolution reveals a cooccurrence of RNA modifications across the transcriptome"

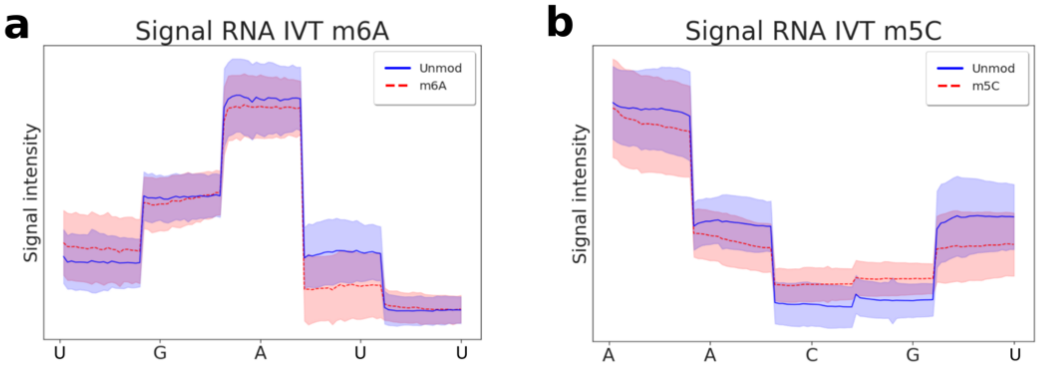


**Supplementary Figure 1. Nanopore DRS signals for sites harboring modified and unmodified nucleotides. (a)** Characteristic profiles of multiple read signals from *in vitro* transcripts (IVTs) with N6-methyladenosine (m6A) (in red) or adenosine (in blue) (Unmod) located in the middle position. Shown are nanopore current signals from five consecutive nanopore translocation events, i.e., five consecutive overlapping 5-mers forming a 9-mer centered around the nucleotide of interest. The signal from each overlapping 5-mer is scaled over the central nucleotide of that 5-mer to twenty values and plotted atop the position of its central nucleotide. Only the central nucleotide of each 5-mer is shown on the x-axis. The solid lines show the mean values, and the shades indicate the 95% confidence intervals of the mean. **(b)** Same as (a), but for the read signals from IVTs containing 5-methylcytidine (m5C) (in red) and cytidine (in blue) (Unmod).

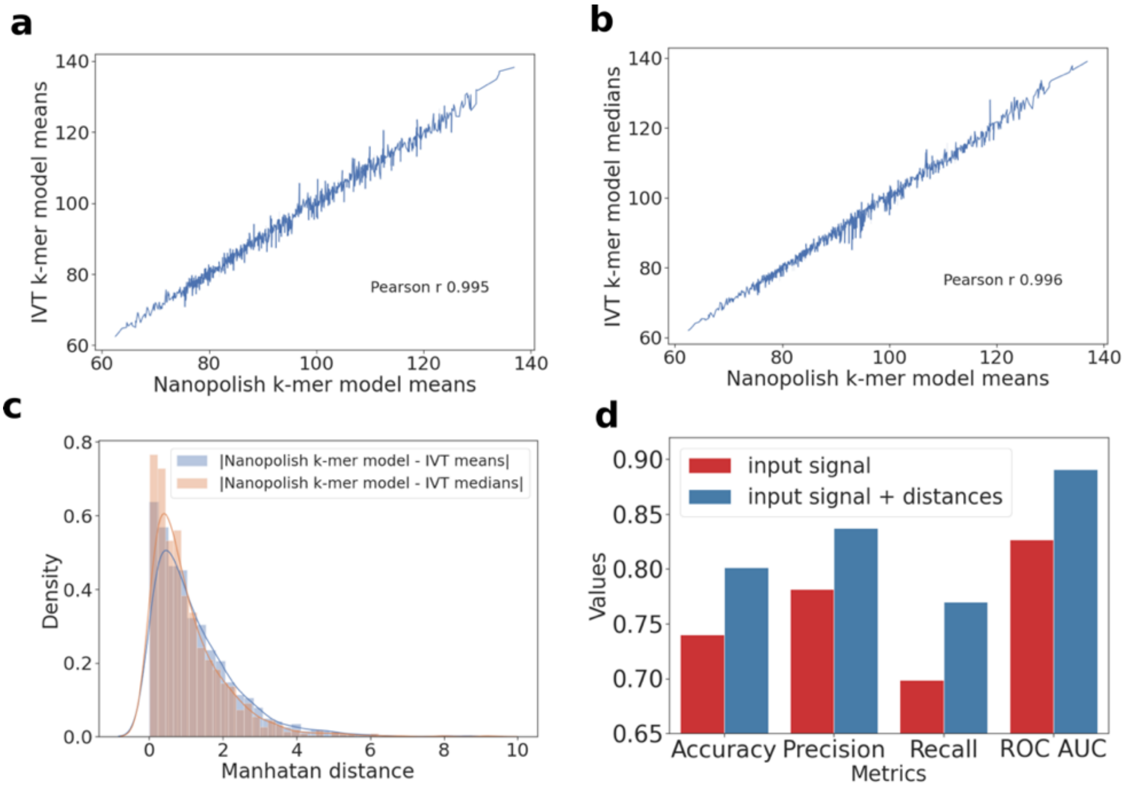


**Supplementary Figure 2. Properties of the nanopore current signals and the effect of a distance vector in CHEUI-solo' s performance metrics. (a)** Correlation of the values from the Nanopolish k-mer model (x-axis) (k=5) with the mean signal values calculated for the same k-mers (k=5) using signals from unmodified *in vitro* transcripts (IVTs) (y-axis). **(b)** Correlation of the values from the Nanopolish k-mer model (x-axis) (k=5) with the median signal values calculated for the same k-mers (k=5) using signals from unmodified IVTs (y-axis). **(c)** Distribution of the absolute differences (x-axis) between the values estimated from the unmodified IVT signals and those from the Nanopolish k-mer model. The line plot represents the fitting of the histogram to a density function. **(d)** Comparison of the performance of the CHEUI m6A model 1 with (blue) and without (red) using the distance vector as input. As described in the Methods section, the distance vector is calculated as the vector of absolute differences between the observed and Nanopolish k-mer model (k=5) values in each of the 100 components derived from the signals associated with each 9-mer in each read. CHEUI performance is assessed by the accuracy, precision, recall, and area under the receiver operating characteristic (ROC) curve (AUC) from the IVT test 1 set (x-axis).

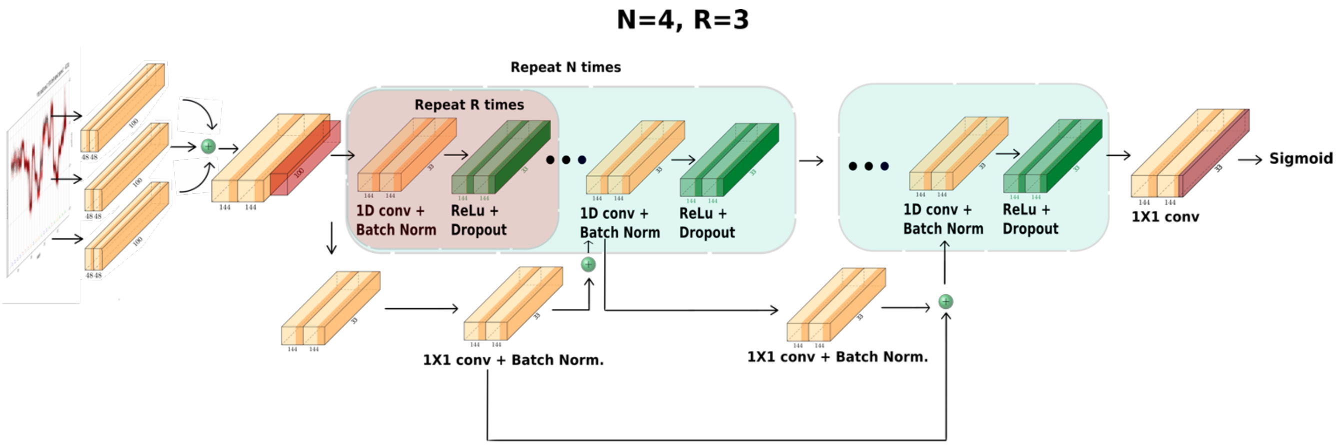


**Supplementary Figure 3. Schematic view of CHEUI-solo convolutional neural network models.** Both Model 1 and Model 2 in CHEUI-solo are convolutional neural networks (CNNs) defined as shown in the diagram and are based on the CNN architecture employed in fast speech recognition algorithms[1]. This architecture is defined by N blocks and R sub-blocks. For CHEUI we used 4 blocks (green) (N=4 in the diagram), each containing 3 sub-blocks (red) (R=3 in the diagram). Sub-blocks are made of 1D convolutions (1D conv.), followed by batch normalization (Batch Norm.), a Rectified Linear Unit (ReLu), and a dropout. All sub-blocks in a block have the same number of output channels. The filter size is 2 for the first block and 3 for the rest of the blocks. The number of filters was set to 48 for all the blocks. Each block input is connected directly into the last sub-block via a residual connection. The residual connection is first projected through a 1×1 convolution to account for different numbers of input and output channels, then through a batch norm layer. The output of this batch norm layer is added to the output of the batch norm layer in the last sub-block. The result of this sum is passed through the activation function and dropout to produce the output of the current block. Both CNNs were implemented using Keras[2] and Tensorflow[3]. However, the inputs for each CNN were different. For CHEUI-solo Model 1, we used two vectors as inputs: the observed signal values and the difference with the expected signal values, for each 9-mer. CHEUI-solo Model 2 uses a single vector describing the distribution of the CHEUI-solo Model 1 probability for the reads at a given transcriptomic site. In both models, the classification is binary, i.e., modified, or unmodified.

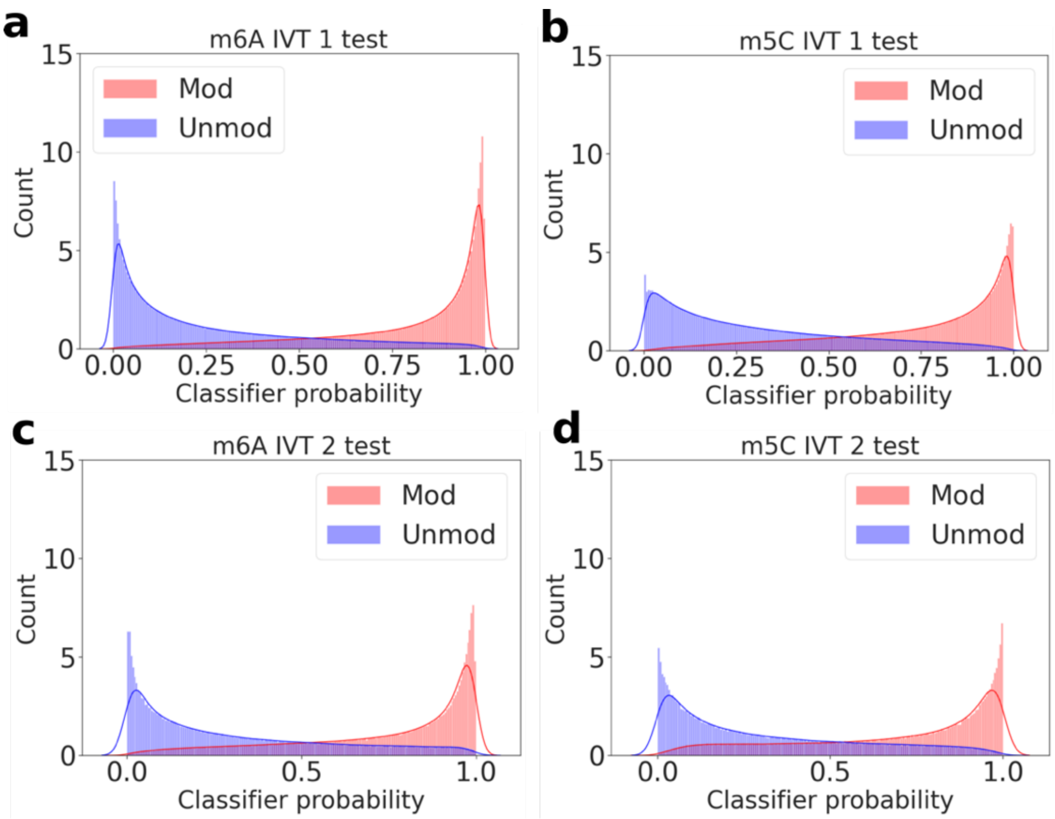


**Supplementary Figure 4. Distribution of CHEUI's individual read probabilities.** Shown are probability distributions returned by CHEUI-solo Model 1 for all the sites and all the reads in the IVT test 1 dataset, for the positive (Mod) and negative (Unmod) cases for m6A **(a)** and m5C **(b)**. **(c, d)** same as (a b), but for the IVT test 2 dataset.

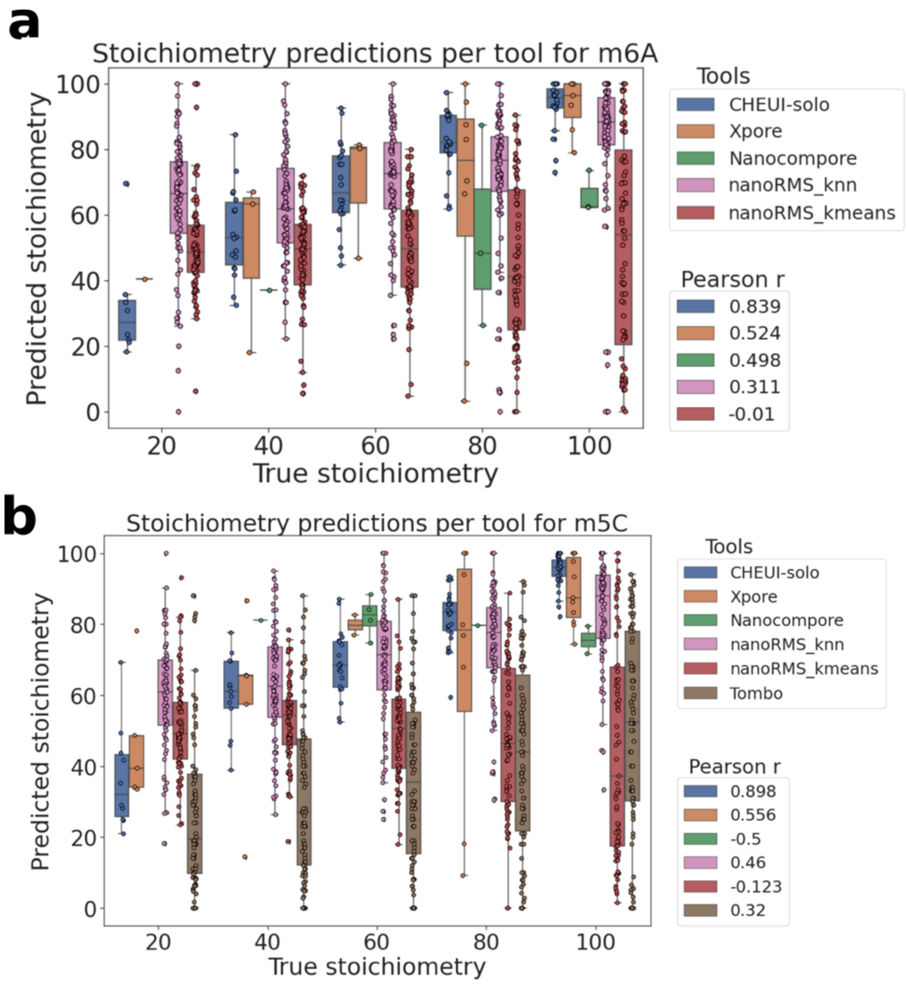


**Supplementary Figure 5. Benchmarking the modification stoichiometry prediction by CHEUI in comparison to other available tools.** We show the stoichiometry values predicted by each tool (y-axis) and the actual stoichiometry values of the controlled read mixtures (x-axis) for m6A **(a)** and for m5C **(b)**. Tool names and Person r-values for the correlation with the ground truth stoichiometry are given in the legends on the right.

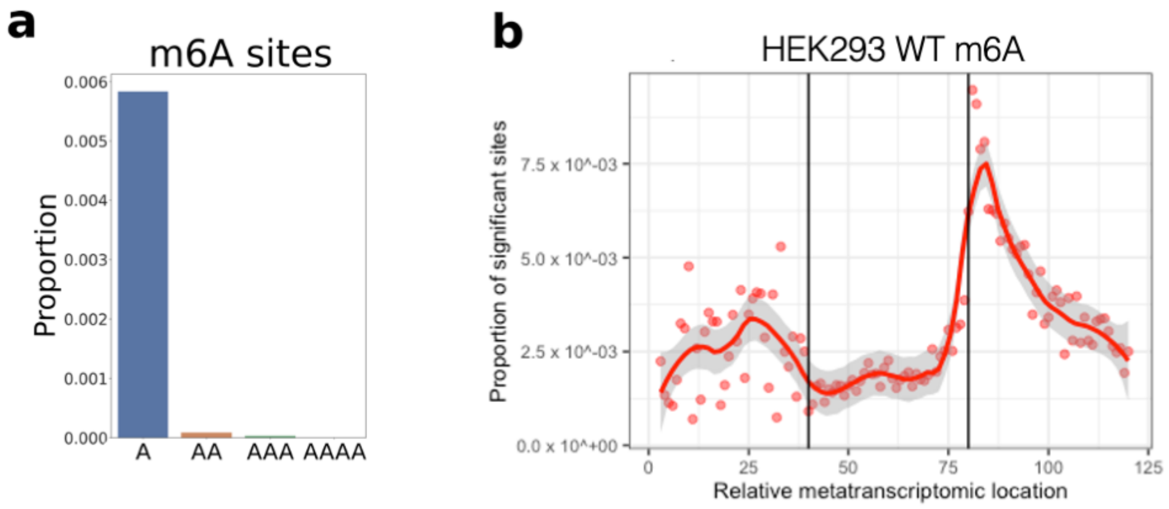


**Supplementary Figure 6. CHEUI m6A predictions in HEK293 WT cells. (a)** We show the proportion of sites of the form A, AA, AAA or AAAA, where all A positions were predicted as m6A. **(b)** We show the proportion of significant transcriptomic sites over tested sites on mRNAs. Sites are mapped and rescaled into three equal-sized segments (each defined to be 40 positions) according to the region in the mRNA: 5′ untranslated region (UTR), coding sequence (CDS), and 3′ UTR. The proportions are shown for the WT sample. Solid lines indicate a LOESS fit to the proportions along the x-axis (represented as dots), and the shades indicate the 95% confidence intervals of the LOESS fit.

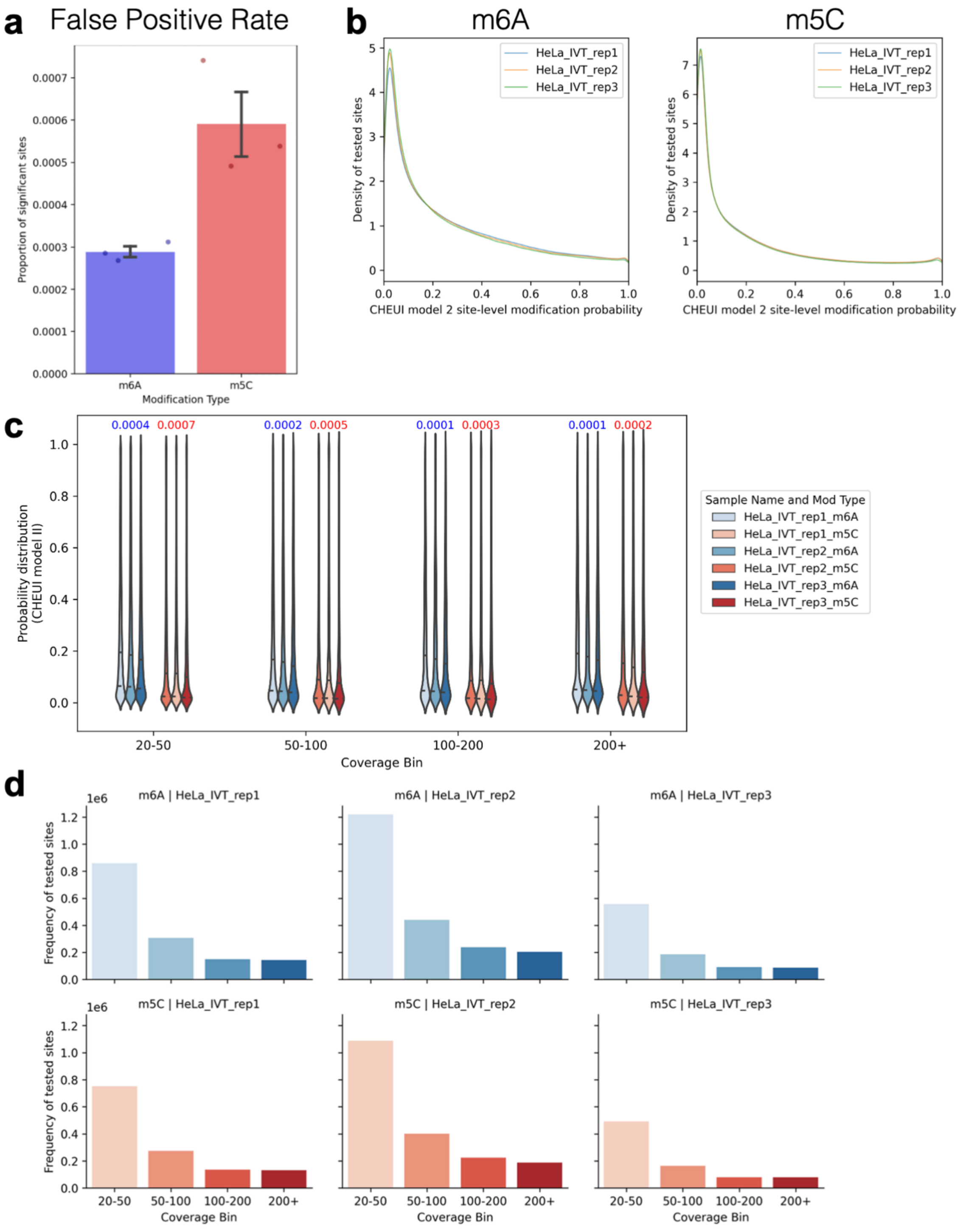


**Supplementary Figure 7. Analysis of an *in vitro* transcribed transcriptome with CHEUI. (a)** The plot shows the proportion of sites predicted (CHEUI Model 2 probability > 0.9999) for m6A (left) and m5C (right) in each of the three replicates of DRS from *in vitro* transcribed (IVT) WT HeLa cells[5]. **(b)** The density plots for the CHEUI Model 2 probability for all sites tested for m6A (left and m5C (right). **(c)** Distribution of the CHEUI Model 2 probability values (y axis) for different ranges of read coverage (x axis) for each replicate, for m6A and m5C. The numbers at the top correspond to the average proportions of false predictions in the given coverage range across the three replicates. **(d)** Number of tested sites (y axis) for each of the coverage ranges (x axis) in each HeLa IVT replicated for the m6A (upper panels) and m5C (lower panels) models.

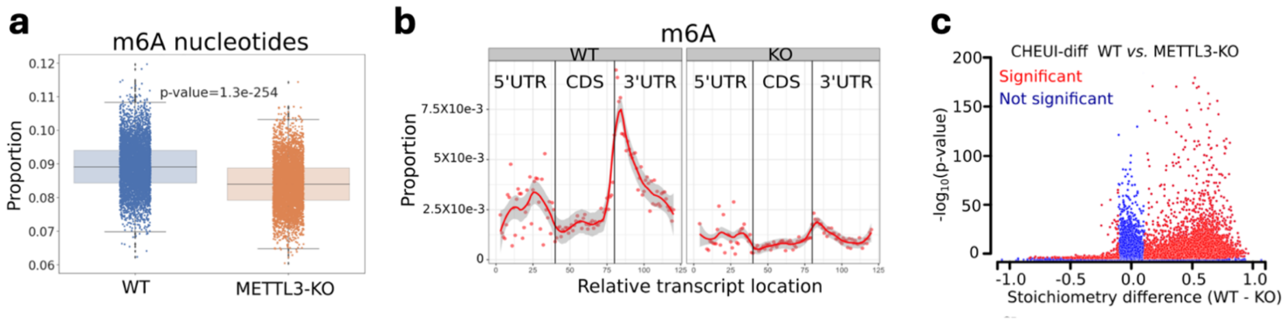


**Supplementary Figure 8. CHEUI detects reduction of m6A in METTL3-KO cells. (a)** Boxplots showing the proportions of modified A (m6A) over the total of A nucleotides (modified and unmodified) in the WT and METTL3-KO samples. Each dot indicates the proportion of individual nucleotides in individual reads predicted as modified by CHEUI-solo Model 1 (probability > 0.9), from a random pool of 5,000 individual nucleotides tested. The box plot represents the four quartiles of the distribution built from 5,000 random samples over those 5,000 nucleotides. The shaded boxes indicating the second and third quartiles, and the bottom and top bars representing the first and fourth quartiles. A non-parametric one-tailed rank-sum test was used to test for statistical significance. **(b)** We show the proportion of significant transcriptomic sites over tested sites on mRNAs. Sites are mapped and rescaled into three equal-sized segments (each defined to be 40 positions) according to the region in the mRNA: 5′ untranslated region (UTR), coding sequence (CDS), and 3′ UTR. The proportions are shown for the WT sample (left panel) and the METTL3-KO sample (right panel). Solid lines indicate a LOESS fit to the proportions along the x-axis (represented as dots), and the shades indicate the 95% confidence intervals of the LOESS fit. **(c)** Results from CHEUI-diff comparing 3 WT and 3 METTL3-KO replicates. Every dot represents a transcriptomic site, with its significance given as $-\log_{10}$(p-value) (y-axis) and the difference in the stoichiometry between WT and METTL3-KO (x-axis).

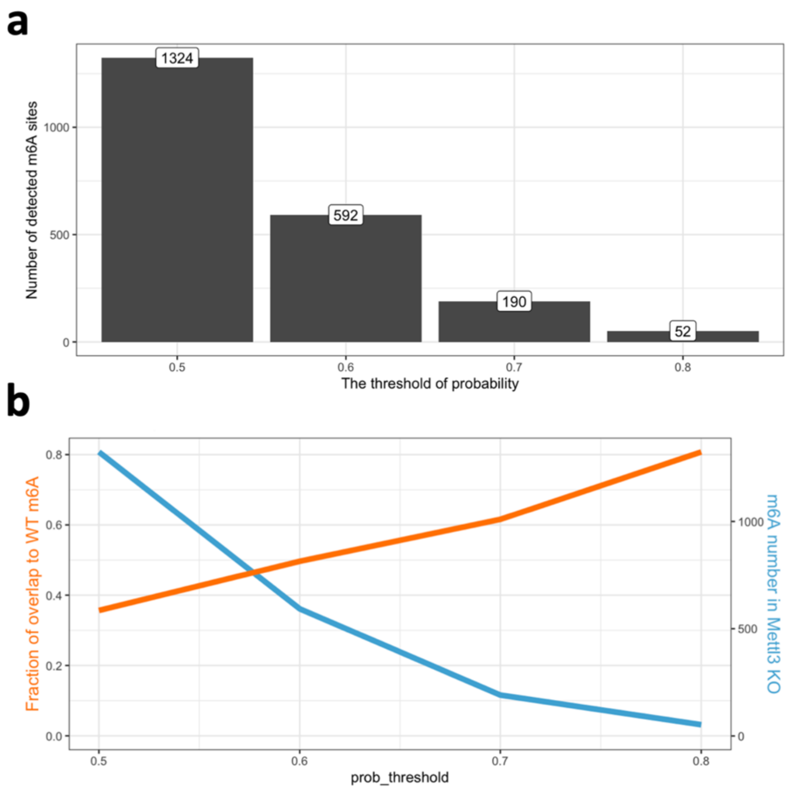


**Supplementary Figure 9.** We used m6Aboost[4] (https://github.com/ZarnackGroup/m6Aboost), an independent machine-learning method that processes miCLIP2 features to predict m6A. We could identify up to 1,324 m6A sites in a Mettl3 KO in mouse embryonic stem cells. (a) The number of m6A sites in the Mettl3 KO (y-axis) at different thresholds (x-axis). (b) Overlap between the Mettl3 KO and WT m6A sites (y-axis) at different thresholds (x-axis).

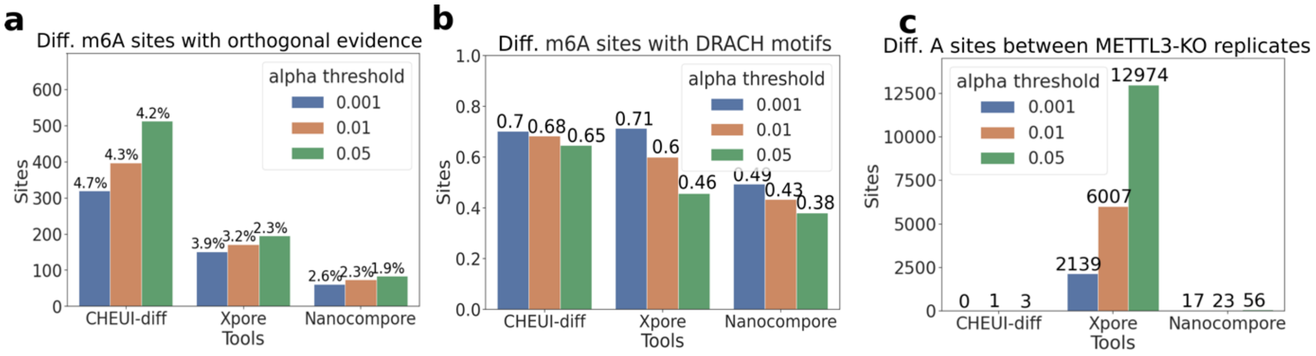


**Supplementary Figure 10. Analysis of the differential modification site detection and quantification in METTL3-KO *vs.* wild-type cells by CHEUI in comparison with the other available tools. (a)** The number of differentially modified m6A sites between HEK293 WT and METTL3-KO predicted by CHEUI-diff, Xpore, and Nanocompore at three levels of significance and overlapping the experimental site set from m6ACE-seq or miCLIP obtained for HEK293. The percentages on the top of the bars correspond to the percentage of sites with orthogonal evidence with respect to the total number of predicted differential sites. **(b)** The proportion of differential significant m6A sites containing a DRACH motif for each method at three levels of significance. **(c)** Differential m6A significant sites discovered by CHEUI, Xpore, and Nanocompore comparing two different METTL3-KO biological replicates.

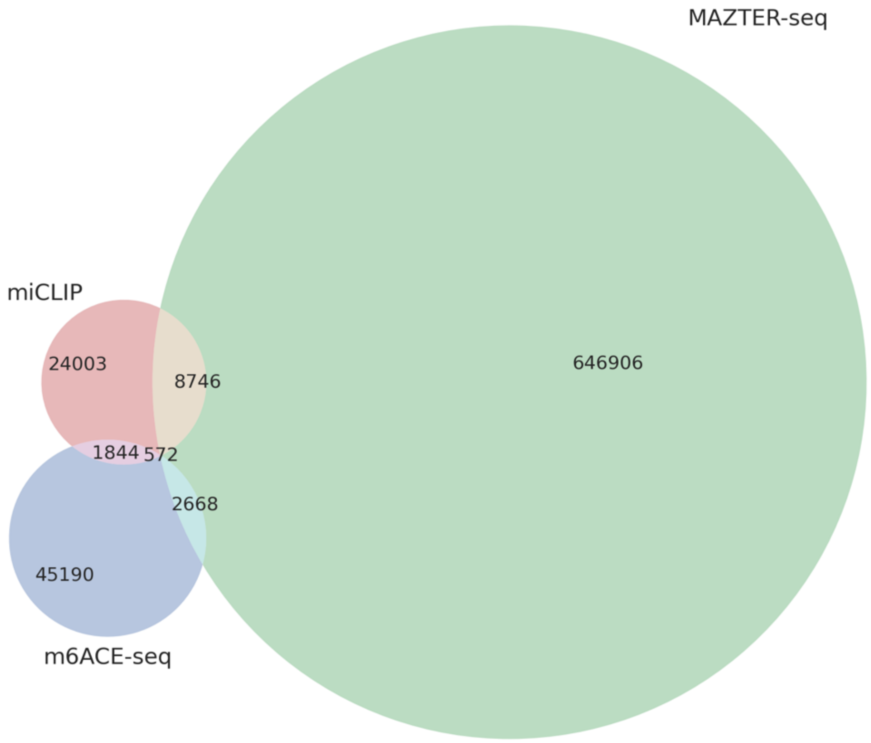


**Supplementary Figure 11. Overlap between m6A sites identified by different experimental techniques.** We show the overlap between the sites identified by m6ACE-seq[6], miCLIP-seq[7], and MAZTER-seq[8] in HEK293 cells. The sites published in each case were downloaded and tested for position overlap. The differences in the same cell model suggest a different coverage between technologies for genes and transcripts, differences in detection limits, and additional possible effects due to the differences in thresholds and detection power.

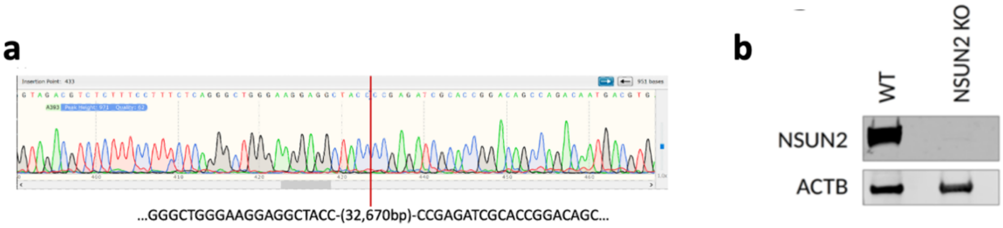


**Supplementary Figure 12. Validation of the NSUN2-KO HeLa cell line. (a)** The edited individual clones were sorted twice. The selected clones were expanded and genotyped using long-range PCR and Sanger sequencing to verify their purity and determine the genomic deletion size. The Sanger sequencing trace shown in the figure confirmed a clean homozygous deletion of 32,670 bp. **(b)** Western blot with an anti-NSUN2 antibody was performed using cell lysates of WT and NSUN2-KO HeLa cells. The signal from actin B antibody (ACTB) is shown for each condition as a loading control.

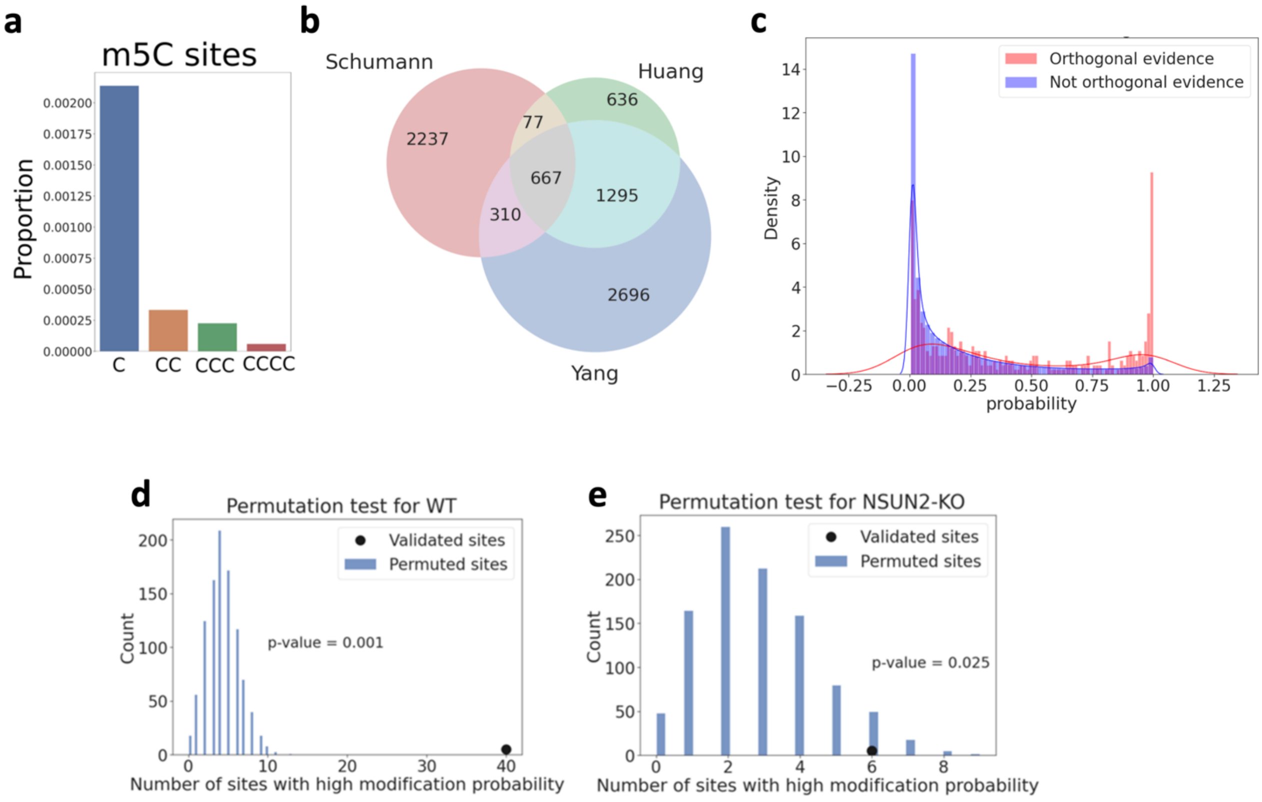


**Supplementary Figure 13. Detection of m5C occurrence in the RNA of HeLa cells with CHEUI. (a)** We show the proportion of sites C, CC, CCC, or CCCC, where all C positions were predicted as m5C in WT HeLa. **(b)** Overlap between the transcriptome m5C sites in HeLa obtained from three different bsRNA-seq experiments Schumann *et al.* 2020 (Schuman)[11], Huang *et al.* 2019 (Huang)[12], and Yang *et al.* 2017 (Yang)[13]. **(c)** Distributions of CHEUI modification detection probability from CHEUI-solo Model 2 for all tested m5C sites before the probability cutoff >0.9999 (x-axis), split according to whether they had orthogonal evidence (red) or not (blue) from the bisulfite RNA sequencing (bsRNA-seq). We only used bsRNA-seq sites that had at least 20 reads in the DRS data to be able to assign a CHEUI-solo Model 2 probability. In **(d)** and **(e)** we compare CHEUI m5C detections in WT and NSUN2-KO HeLa cells with calls derived from bsRNA-seq. Bisulfite detection-based m5C sites are enriched in high CHEUI probabilities. The number of transcriptomic m5C sites predicted by bisulfite sequencing in HeLa cells that had a CHEUI-solo Model 2 probability > 0.99 (black dot) is shown in comparison to the number of cases obtained by chance (blue bars). The cases obtained by chance were calculated by shuffling the CHEUI probabilities across tested sites 10,000 times and calculating again each time how many of the bisulfite-based m5C sites had a probability > 0.99. Shown are the results using Nanopore signals from the WT **(d)** and the NSUN2-KO **(e)** HeLa cells.

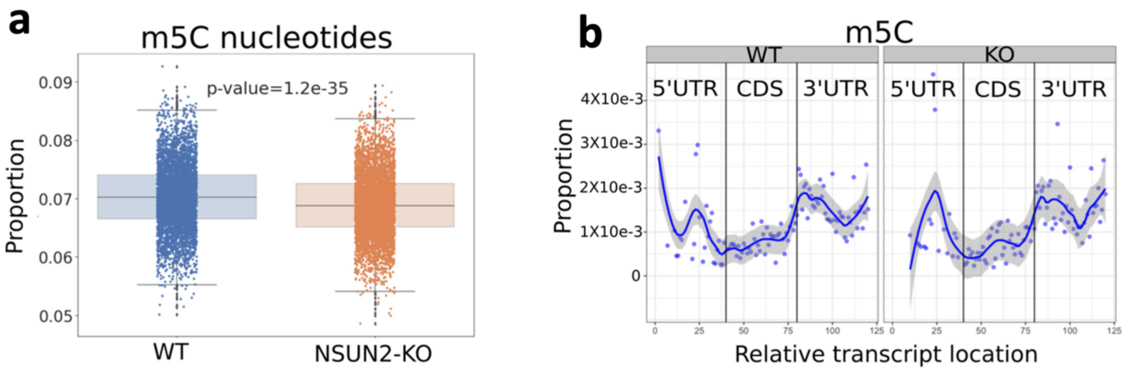


**Supplementary Figure 14. CHEUI detects a mild reduction of m5C in NSUN2-KO HeLa cells. (a)** Boxplots showing the proportions of modified C nucleotides (m5C) over the total of C nucleotides (modified and unmodified) in the WT and NSUN2-KO HeLa samples. Each dot indicates the proportion of individual nucleotides in individual reads predicted as modified by CHEUI-solo Model 1 with probability > 0.9 from a random pool of 5,000 individual nucleotides tested. The box plot represents the four quartiles of a distribution built from 5,000 random samples of those 5,000 nucleotides. The shaded boxes indicate the second and third quartiles, and the bottom and top bars represent the first and fourth quartiles. Non-parametric one-tailed rank sum test was used to test for the statistical significance. **(b)** We show the proportion of significant m5C transcriptomic sites over the total of tested sites on mRNAs. Sites are mapped and rescaled into three equal-sized segments (each defined to be 40 positions) according to the region in the mRNA: 5′ untranslated region (UTR), coding sequence (CDS), and 3′ UTR. The proportions are shown for the WT sample (left panel) and the NSUN2-KO sample (right panel). Solid lines indicate a LOESS fit to the proportions along the x-axis (represented as dots), and the shades indicate the 95% confidence intervals of the LOESS fit.

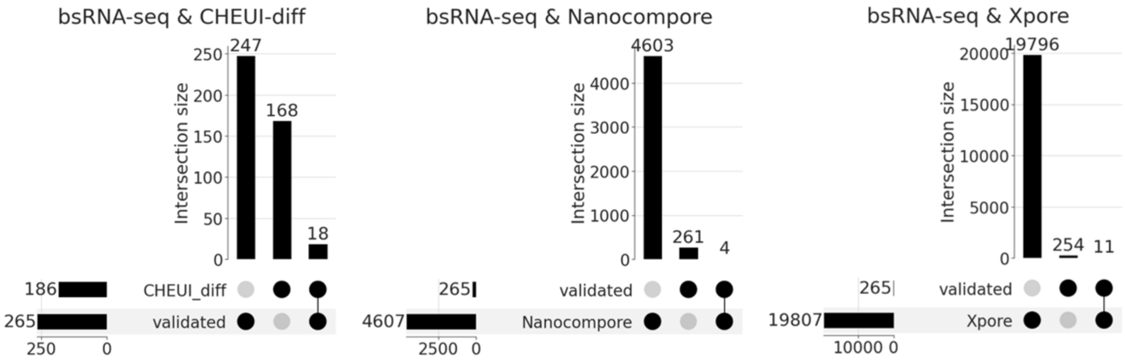


**Supplementary Figure 15. Comparison of nanopore-based m5C detection with bisulfite RNA sequencing.** We show the overlaps between CHEUI-diff (left), Nanocompore (middle), and Xpore (right) with m5C sites in HeLa obtained from three different bisulfide RNA sequencing (bsRNA-seq) experiments Schumann *et al.* 2020 (Schuman)[11], Huang *et al.* 2019 (Huang)[12], and Yang *et al.* 2017 (Yang)[13]. CHEUI, Nanocompore, and Xpore were used to detect the sites that had a significant reduction of stoichiometry when comparing HeLa WT and HeLa NSUN2-KO. The bsRNA-seq sites are those detected in at least one of the three experiments. The plots show the number of detected sites by each method (horizontal bars) and the number of sites that are predicted by either method or both (vertical bars).

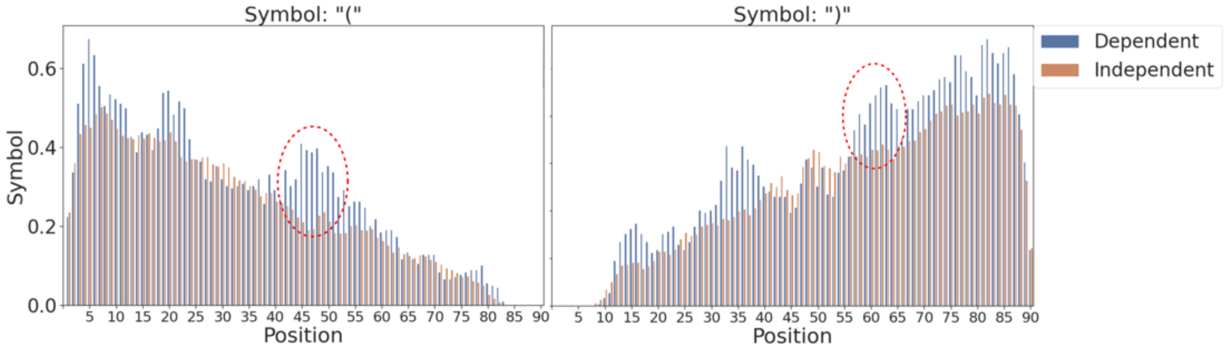


**Supplementary Figure 16. Overview of predicted RNA secondary structure formation propensity associated with the m5C sites detected by CHEUI.** Shown is the proportion of paired bases, indicated with the symbol '(' (left panel) or ')' (right panel) in the predicted secondary structures using 90 nt around (45 nt on either side) the predicted m5C sites on transcriptomic sequences, removing sites that appeared in more than one transcript in the same gene. Position 45 contains the m5C predicted by CHEUI-solo **(a)**. The dashed red circle indicates positions that demonstrate an increase of base-paired nucleotides potentially forming a stem-loop at sequences from NSUN2-dependent sites.

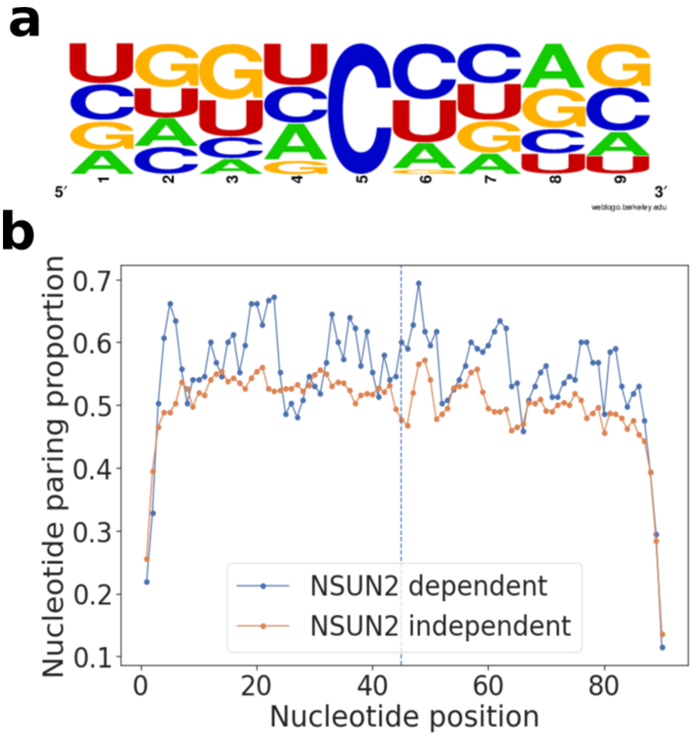


**Supplementary Figure 17. Sequence motifs and RNA secondary structure predictions in m5C sites. (a)** Sequence motifs for NSUN2-independent sites defined as significant sites by CHEUI-solo in the NSUN2-KO sample (1,841 transcriptomic sites). **(b)** Proportion of base-paired positions along 90 nt centered at the m5C sites (as in **Fig. 4h** of the main text) predicted by CHEUI-solo. The vertical line indicates the m5C position. NSUN2-dependent sites were defined as sites predicted by CHEUI-diff to have a significant decrease in modification stoichiometry in the NSUN2-KO sample, whereas NSUN2-independent sites were defined in this case as sites significant according to CHEUI-solo for the NSUN2-KO sample. Regarding their genomic positions, we had 1,250 m5C sites in HeLa WT that do not change in the HeLa NSUN2-KO (definition 1), and 1,940 m5C sites in HeLa NSUN2-KO (definition 2), which we considered as alternative definitions of NSUN2-indendent sites. From those sites, about 43% (833 of 1,940) of them in definition 2 coincide with sites in definition 1, and about 66% (833 of 1,250) of sites from definition 1 coincide with sites from definition 2. While there is a considerable overlap that may explain part of the similarity of the two approaches, there are many other sites that independently contribute to this result.

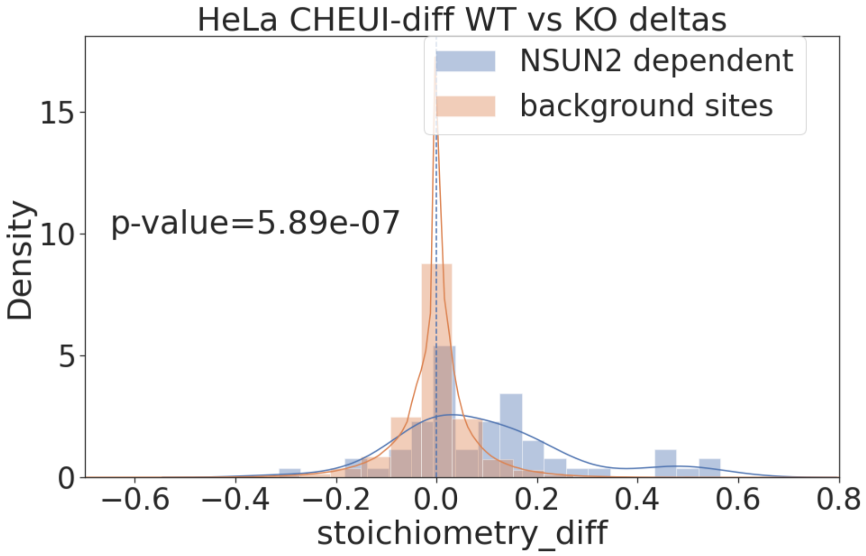


**Supplementary Figure 18. Distribution of the stoichiometry difference reported by CHEUI-diff in NSUN2-dependent and background sites.** Blue bars indicate the stoichiometry difference between WT and NSUN2-KO samples in sites that were reported previously to be NSUN2-dependent by Huang *et al.* 2019[12]. Orange bars indicate the stoichiometry difference between WT and NSUN2-KO sites that were not in the NSUN2-dependent list (i.e., not significant according to CHEUI-diff) and were not found on the list of Huang *et al.* 2019. A non-parametric one-tailed test using rank-sum was performed to compare the two distributions.

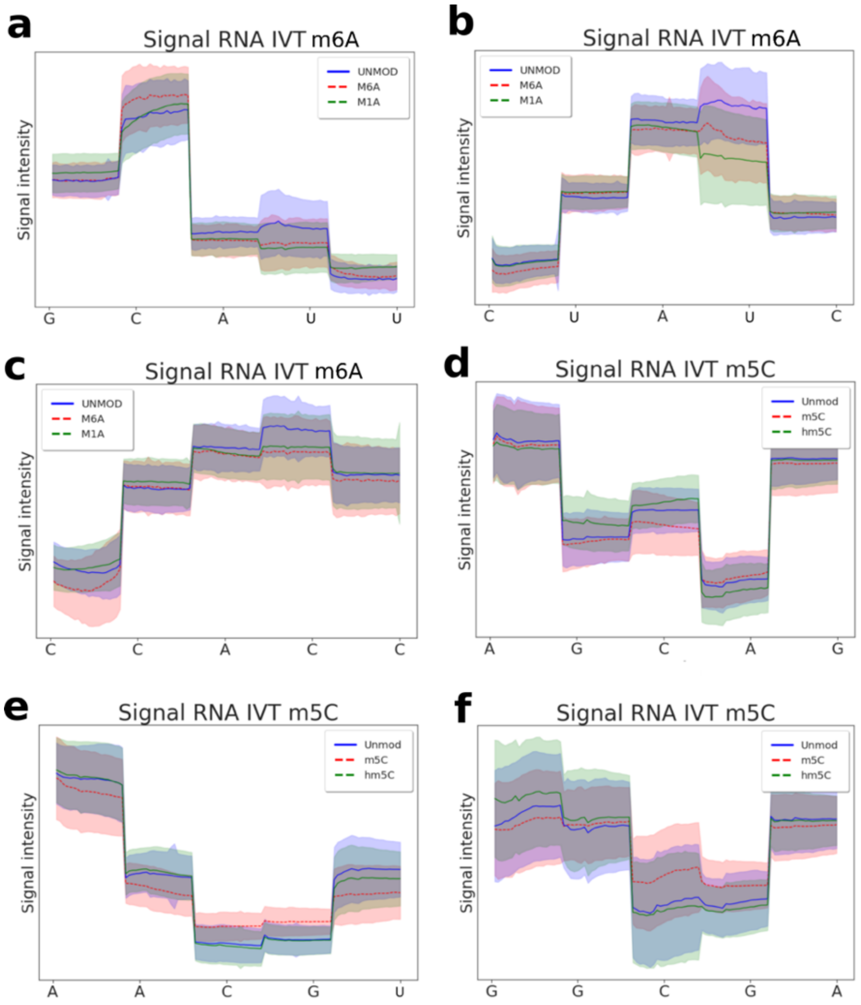


**Supplementary Figure 19. Visualization of nanopore current signals for sites harbouring different A and C modifications against non-modified sites.** Shown are nanopore current signals from five consecutive nanopore translocation events, i.e., five overlapping 5-mers forming a fixed 9-mer. The signal from each overlapping 5-mer is scaled over the central nucleotide of that 5-mer to twenty values and plotted atop the position of its central nucleotide. Only the central nucleotide of each 5-mer is shown on the x-axis. The line shows the mean values, and the shades indicate the 95% confidence intervals of the mean. Three different sequence contexts are shown exemplifying typical signal profiles for A (Unmod), m6A, and m1A (**a**, **b**, **c);** and for C (Unmod), m5C, and hm5C **(d, e, f)**.

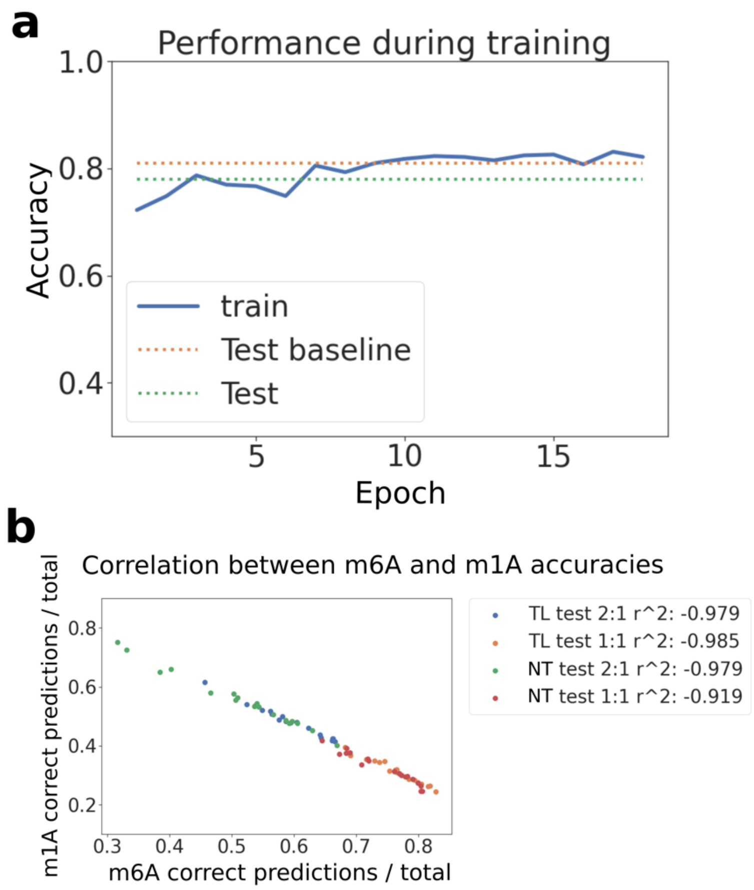


**Supplementary Figure 20. Accuracy metrics returned by CHEUI-solo Model 1 upon retraining with other modifications as negatives.** We retrained CHEUI-solo using the signals for m1A and all other modifications different from m6A as part of the negative set for the training. **(a)** Accuracy (y-axis) of CHEUI Model 1 retrained to predict m6A using other available RNA modifications read signals (Y, f5C, m7G, m1A, I, hm5C, m5C) plus unmodified signals as negatives, and using the m6A read signals from IVT train 1 as positives. Accuracy was calculated using m6A and unmodified signals during training ('train') and on the IVT test 1 dataset ('Test'), compared to the original CHEUI-solo Model 1 accuracy on IVT test 1 for m6A ('Test baseline'). **(b)** Association between the correctly classified m1A nucleotides divided by the total number of m1A nucleotides (y-axis) and the correctly classified m6A nucleotides divided by the total of m6A nucleotides (x-axis) for different versions of CHEUI-solo Model 1, using transfer learning from the original model (TL) or not (DL) and using different label weights, i.e., relative contribution of the negative examples in the loss function (1:1, 2:1).

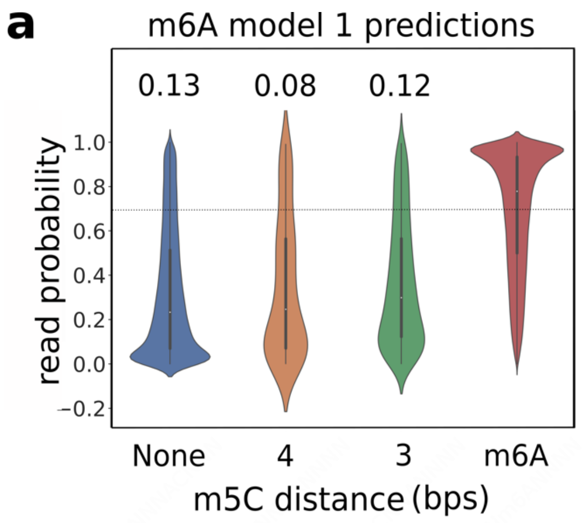


**Supplementary Figure 21. CHEUI's prediction probability of m6A at individual read level.** IVT2 read signals at 9-mers were centered at A (NNNNANNNN) with various configurations: 9-mers with no m6A ('None'), 9-mers with m5C present 3 ('3') or 4 ('4') nucleotides from the central A, and 9-mers with a modified middle A ('m6A'). We could not find any 9-mers NNNNANNNN in the IVT test 2 dataset with only one m5C at distances 1 or 2 nucleotides from the middle A.

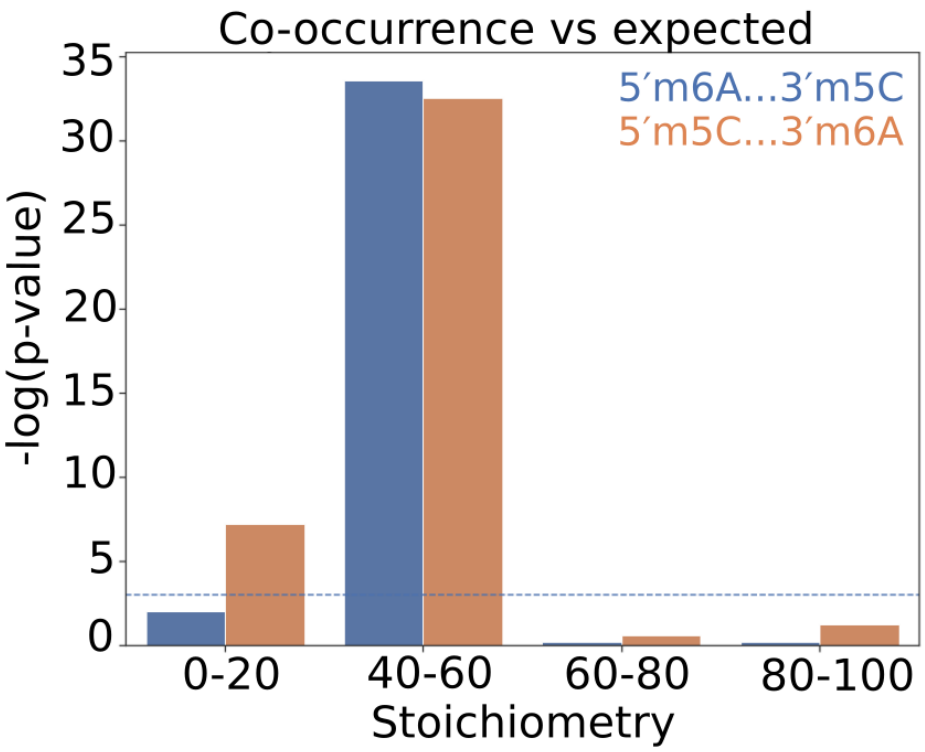


**Supplementary Figure 22. Co-occurrence of m6A and m5C at different stoichiometry values.** We grouped pairs of CHEUI-solo m6A-m5C and m5C-m6A sites in the same transcript according to their mean stoichiometry and compared their co-occurrence values in reads with the expected ones. The expected values were obtained by permuting the reads on each site individually and controlling for the stoichiometry in the permutations. We plot the significance of the co-occurrence across individual reads (y-axis) as a function of the average stoichiometry of both sites (x-axis). The Mann-Whitney U-tests between co-occurrence and permutated values were performed at different stoichiometry levels. The pairs of m6A and m5C sites were collected when falling between 5 to 30 nucleotides away from each other.

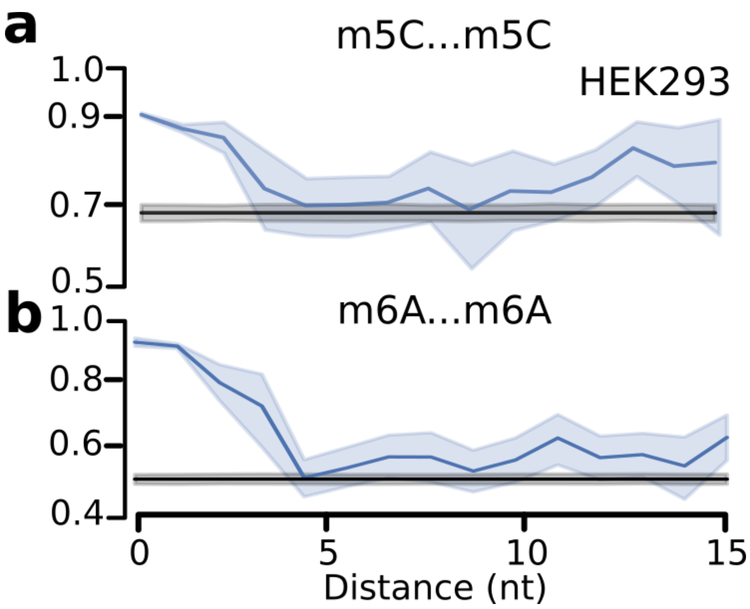


**Supplementary Figure 23. Read-level co-occurrence of the same modification. (a)** Read-level co-occurrence of m5C-m5C at various relative distances (x-axis). Co-occurrence is calculated by counting the number of reads where two sites have the same modification state, either both modified (Model 1 probability > 0.7) or both unmodified (Model 1 probability < 0.3) divided by the total number of reads covering both sites. The black line and grey shades indicate the mean and 95% confidence interval values for the expected co-occurrence values, obtained by permuting the modification status of reads at each site individually. Distances are measured as the difference between positions of the two modified nucleotides, e.g., 5′-m6ANNNNm5C-3′ are at the relative distance of 5 nt. **(b)** Same as (a) but for the co-occurrence of m6A-m6A.

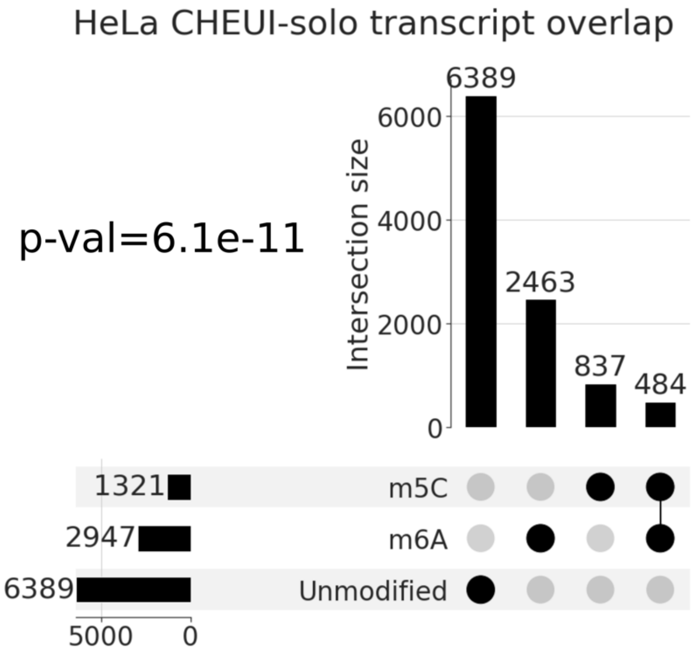


**Supplementary Figure 24. CHEUI-solo identifies co-occurrence of m6A and m5C sites on mRNAs in HeLa.** UpSet[14] plot for the number of reference transcripts containing m6A and m5C sites, transcripts containing only one of the modifications, and transcripts containing none, as identified by CHEUI-solo. The Fisher's test p-value shows the association of m6A and m5C sites relative to all the tested sites. Only cases where m6A and m5C are at a relative distance of 5 nt or more were considered.

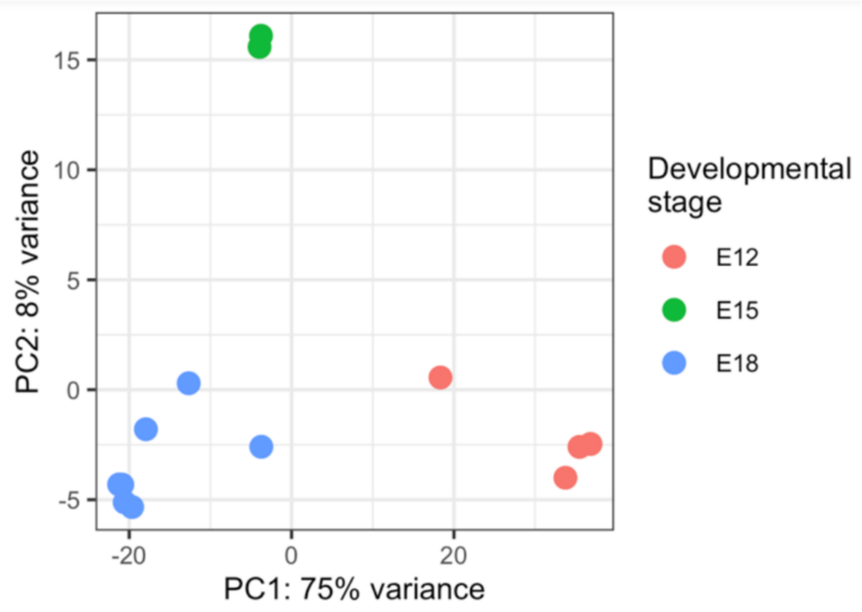


**Supplementary Figure 25. Nanopore DRS in mouse pre-frontal cortex.** Principal Component Analysis (PCA) plot of the replicate samples sequenced at embryonic development stages E12, E15, and E18. The PCA plot was generated using log-normalized gene counts from the DESeq2 output.

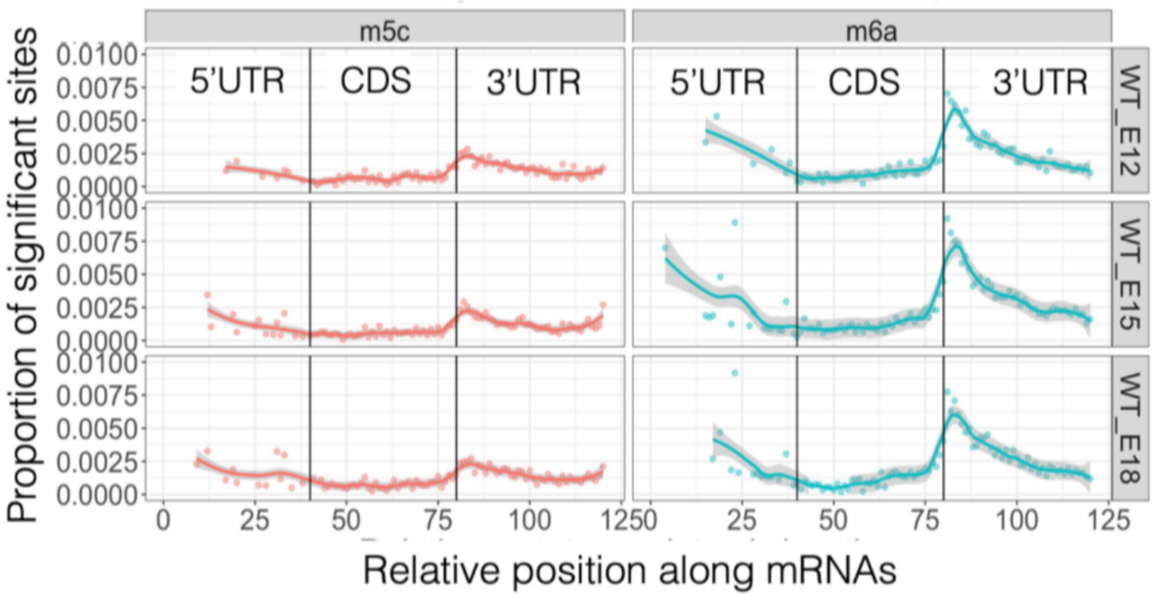


**Supplementary Figure 26. Methylation profiles of mRNAs across the three stages of mouse embryonic pre-frontal cortex development reported by CHEUI-solo.** Shown are the proportion of significant m5C (left panels) and m6A (right panels) transcriptomic sites over the total of tested sites on mRNAs. Sites are separated into three equal-sized segments (of length 40 nt) according to the region in the mRNA: 5′ untranslated region (UTR), coding sequence (CDS), and 3′ UTR. The proportions are shown for the E12, E15 and E18 pre-frontal embryonic mouse samples. Solid lines indicate a LOESS fit to the proportions along the x-axis (represented as dots), and the shades indicate the 95% confidence intervals of the LOESS.

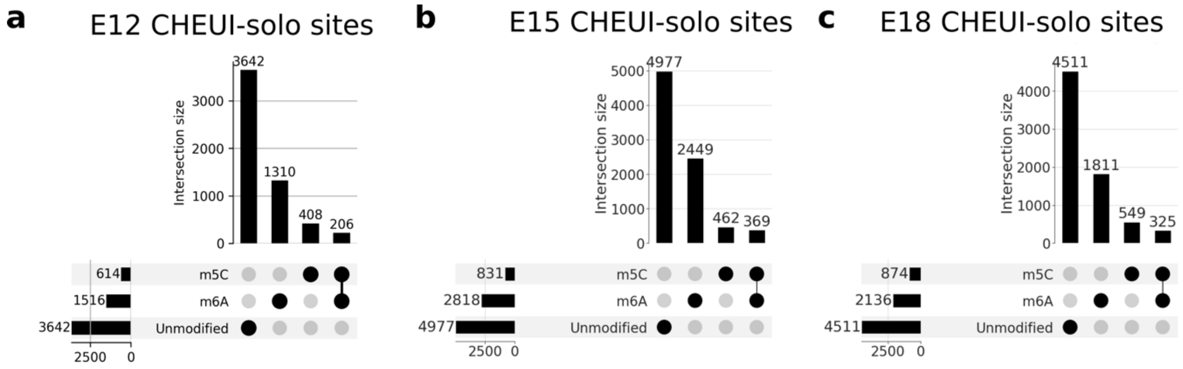


**Supplementary Figure 27. CHEUI-solo identifies co-occurrence of m6A and m5C sites on mRNAs in mouse embryonic pre-frontal cortex.** UpSet[14] plot for the number of protein-coding transcripts from the mouse Ensembl v104 (GRCm39) annotation containing m6A and m5C sites, for each one of the modifications or none of the modification, across the three studied embryonic stages, E12, E15, and E18. Only cases with m6A and m5C at distances of 5 nt or more were considered. In all three cases, the observed co-occurrence of m6A and m5C was significantly higher than expected (Fisher's exact 2-sided test p-value < 0.000150).

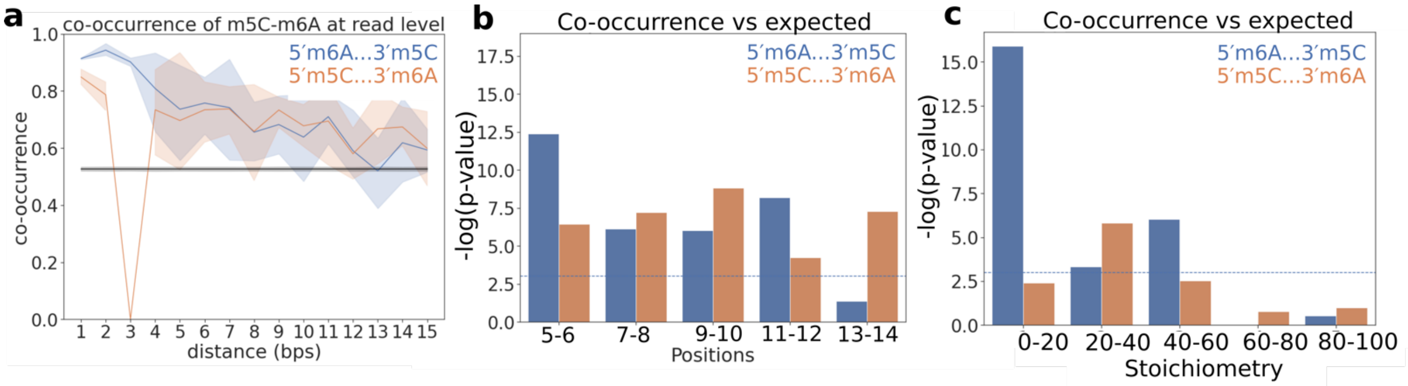


**Supplementary Figure 28. Read-level co-occurrences of m6A and m5C in the RNA of mouse prefrontal cortex.** Data from the 3 development stages (E12, E15, and E18) were grouped together for this analysis. **(a)** Read-level co-occurrence (y-axis) of m6A and m5C modifications at various relative distances (x-axis). Co-occurrence is calculated by counting the number of reads where two sites have the same modification state, either both modified (Model 1 probability > 0.7) or both unmodified (Model 1 probability < 0.3), divided by the total number of reads covering both sites. Pairs of sites with A upstream of C are depicted in blue and pairs of sites with A downstream of C are shown in orange. Dark blue and orange lines indicate mean co-occurrence values, and the shades are the 95% confidence intervals of the mean at each distance. The black line and grey shades indicate the mean and 95% confidence interval, respectively, for the expected co-occurrence values, obtained by permuting the read modification status in each site individually. Distances are measured as the difference between positions of the two modified nucleotides, e.g., 5′-m6ANNNNm5C-3′ are at the relative distance of 5 nt. **(b)** Statistical significance (y-axis) of the comparison between the observed and expected co-occurrences at different distances between pairs of sites (x-axis). The sites were considered in groups of two consecutive distances. Observed and expected co-occurrences were compared using a one-sided Mann-Whitney U-test. A horizontal blue line indicates p-value = 0.05. The expected co-occurrences were calculated by permuting the modification status of reads in each site independently. **(c)** Significance of the co-occurrence across individual reads (y-axis) as a function of the average stoichiometry of both sites (x-axis). We grouped pairs of CHEUI-solo m6A-m5C sites in the same transcript according to their mean stoichiometry of m6A and m5C and compared their co-occurrence values with the expected ones, obtained by permuting the reads on each site individually and controlling for the stoichiometry in the permutations. The Mann-Whitney U-tests between co-occurrence and permutated values were performed at each stoichiometry range. Only pairs of m6A and m5C sites at relative distances of 5-30 nt were considered.

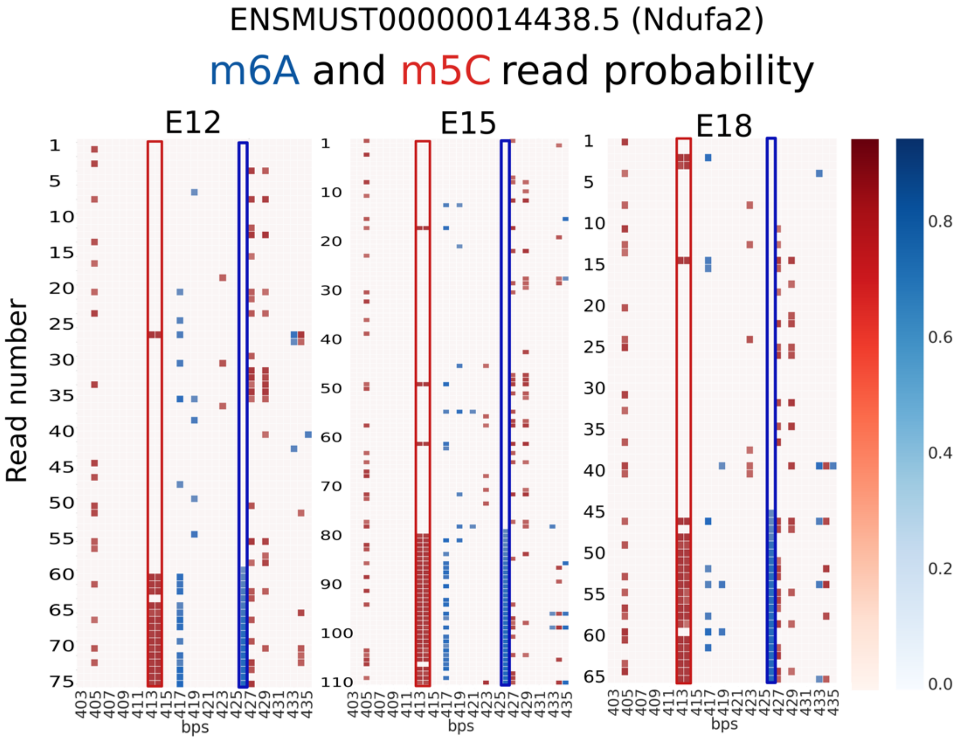


**Supplementary Figure 29. Conservation of m6A and m5C co-occurrence in the RNA of mouse embryonic pre-frontal cortex across developmental stages.** We show m6A and m5C modifications co-occurring in the same RNA reads for transcript ENSMUST00000014438 (gene Ndufa2) at a distance of 13 nt over three mouse embryonic development stages, E12, E15, and E18. Read IDs are represented with numbers on the y-axis and the position along the transcript is represented on the x-axis. The color scales represent CHEUI's detection probabilities at the read level for m6A (in blue) and m5C (in red). Only read-level predictions with a CHEUI-solo Model 1 probability > 0.7 are shown.
